# *DVL1* variants and C-terminal deletions have differential effects on craniofacial development and WNT signaling

**DOI:** 10.1101/2024.02.28.582602

**Authors:** Shruti S. Tophkhane, Sarah J. Gignac, Katherine Fu, Esther M. Verheyen, Joy M. Richman

**Author notes:** Senior author Contact: Joy M. Richman Life Sciences Institute, 2350 Health Sciences Mall, University of British Columbia Vancouver, BC, V6T 1Z3, Canada.

## Abstract

Robinow Syndrome (RS) is a rare disease characterized by craniofacial malformations and limb shortening linked with mutations in seven WNT pathway genes. Our objective was to investigate the functional effects of frameshift mutations the intracellular adaptor protein, Dishevelled (*DVL1*; *c.1519*Δ*T*, p.Trp507Glyfs*142) on chicken craniofacial development. Misexpression of wt (wt) or mutant h*DVL1* variants in vivo caused upper beak shortening (wt*DVL1* n=8/14; *DVL1^1519^*^Δ*T*^ 12/13). At early stages of development, the *DVL1^1519^*^Δ*T*^ inhibited frontonasal mass narrowing, chondrogenesis, and proliferation. To test whether the phenotypes were caused due to the abnormal C-terminal peptide in *DVL1^1519^*^Δ*T*^, we designed two additional constructs. The *DVL1^1519*^* (DVL1^507*^) retains first 30 amino acids of the C-terminus while *DVL1^1431*^* (DVL1^477*^) removes the entire C-terminus. *DVL1^1519*^* injected embryos had normal beaks while *DVL1^1431*^* caused high mortality and the phenotypes were like the *DVL1^1519^*^Δ*T*^. In frontonasal micromass cultures, both *DVL1^1519^*^Δ*T*^ and *DVL1^1431*^* inhibited skeletogenesis while the *DVL1^1519*^* resembled wt*DVL1* and *GFP* cultures. In luciferase assays *DVL1^1519^*^Δ*T*^, *DVL1^1519*^*and *DVL1^1431*^* weakly activated the WNT canonical and non-canonical JNK-PCP pathways compared to wt*DVL1*. Furthermore, we observed that variant DVL1^507*fs^ is stalled in the nucleus similar to hDVL1^477*^, possibly due to the abnormal C-terminus interfering with the nuclear export sequence. wtDVL1 and DVL1^507*^ were distributed in nucleus and the cytoplasm. Our RS-*DVL1*^1519ΔT^ avian model recapitulates the broad face and jaw hypoplasia and demonstrates defects in both branches of WNT signaling. This is the first study to clarify the role of abnormal C-terminus in ADRS and to recognize the importance of an uncharacterized C-terminal sequence.

**Summary Statement:** Functional and biochemical studies on chicken embryos with the Robinow syndrome (RS) *DVL1* variant demonstrate defects in skeletogenesis and both branches of WNT signaling. This is the first study to establish a link between the RS facial defects and the mutated C-terminal sequence. We identified first 30 amino acids of the *DVL1* C-terminus are sufficient for normal development.

## Introduction

Dishevelled1 *(DVL1)*, an intracellular adaptor protein, is a key downstream mediator of canonical and non-canonical WNT pathways (Gao and Chen, 2010). The only genetic disease associated with mutations in *DVL* proteins is ADRS (23/67 patients; MIM#616331, Table S1) (Bunn et al., 2015; Gignac et al., 2023; Hu et al., 2022; White et al., 2015; Zhang et al., 2022). RS cases usually have craniofacial features that are very distinctive including jaw hypoplasia, broad nasal bridge and hypertelorism i.e. wide-set eyes. Patients also have short stature, genital defects, and vertebral defects. The specific the face and limbs phenotypes are unexpected since *Dvl1* is ubiquitously expressed in mouse (Sussman et al., 1994) and chicken (Gray et al., 2009). We were motivated to explore how ADRS mutations in *DVL1* affect such specific aspects of human development.

The Dishevelled (*Dsh*) gene was named after a spontaneous *Drosophila* mutant that had randomized hair and bristle arrangements (Fahmy and Fahmy, 1959). Thereafter, human and mouse homologues were cloned (*DVL1*, *2*, *3).* There are two *DVL* genes in the chicken genome *(DVL1, DVL3)* (Gentzel and Schambony, 2017). The function of all *Dvl* isoforms has been studied in mouse germline knockouts (Wynshaw-Boris, 2012) (Table S2). *Dvl1^-/-^*mice are phenotypically normal but show social interaction abnormalities (Lijam et al., 1997). *Dvl1^-/-^* and *Dvl2^-/-^* mice have severe skeletal malformations, neural tube closure defects, and cochlear defects (Hamblet et al., 2002), while *Dvl1^-/-^*and *Dvl3^-/-^* mice die of unknown cause between E13.5 – E15.5 (Etheridge et al., 2008). There is functional redundancy among the *Dvl* genes since more severe phenotypes are seen in *Dvl* double mutant mice (Gao and Chen, 2010; Gentzel and Schambony, 2017; Hamblet et al., 2002; Lijam et al., 1997). The phenotypes observed in the single or compound *Dvl* knockout mice are not consistent with ADRS.

All DVL protein isoforms contain three domains: N-terminal DIX (Dishevelled and Axin), PDZ (Post-synaptic density protein-95, Disc large tumour suppressor, Zonula occludens-1), and DEP (Dishevelled, Egl-10 and Pleckstrin) domain (Fig 1A,C) (Sharma et al., 2018). In addition, DVL contains a ‘basic region’ between DIX and PDZ domain with serine and threonine residues and a ‘proline-rich region’ between PDZ and DEP domain (Dillman et al., 2013; Mlodzik, 2016; Sharma et al., 2018). The DIX and PDZ domains direct WNT signaling to the β-catenin dependent WNT pathway (canonical) while the PDZ and DEP domains activate the WNT/Planar Cell Polarity pathway (Clevers and Nusse, 2012; Habas and Dawid, 2005; Tauriello et al., 2012). The central PDZ domain participates in both pathways and binds to FZD receptors (Wong et al., 2003). In addition, structure-function studies have shown that the conserved DIX, PDZ, DEP domains are essential for WNT signal transduction (Boligala et al., 2022; Gan et al., 2008; Itoh et al., 2005; Kishida et al., 1999; Qi et al., 2017; Rothbacher et al., 2000; Sharma et al., 2018; Wallingford and Habas, 2005; Weitzman, 2005). The three functional domains of DVL mediate interactions with more than 50 binding partners leading to different signaling outcomes (Sharma et al., 2018). The DEP domain is predominantly involved in recruitment of DVL to the cell membrane. The DEP and DIX domains are necessary for homo-and heterodimerization of DVL followed by subsequent activation of the β-catenin mediated WNT pathway (Ngo et al., 2020). The PDZ domain is essential for physical interaction of DVL with other proteins that function as agonists or antagonists of the WNT pathway and the interaction with the FZD receptors (Sharma et al., 2018). Furthermore, the shuttling of DVL proteins in and out of the nucleus is regulated by the nuclear localization sequence (NLS) located between the PDZ and the DEP domain and a nuclear export sequence (NES) that follows the DEP domain (Fig. 1C).

**Figure 1:**
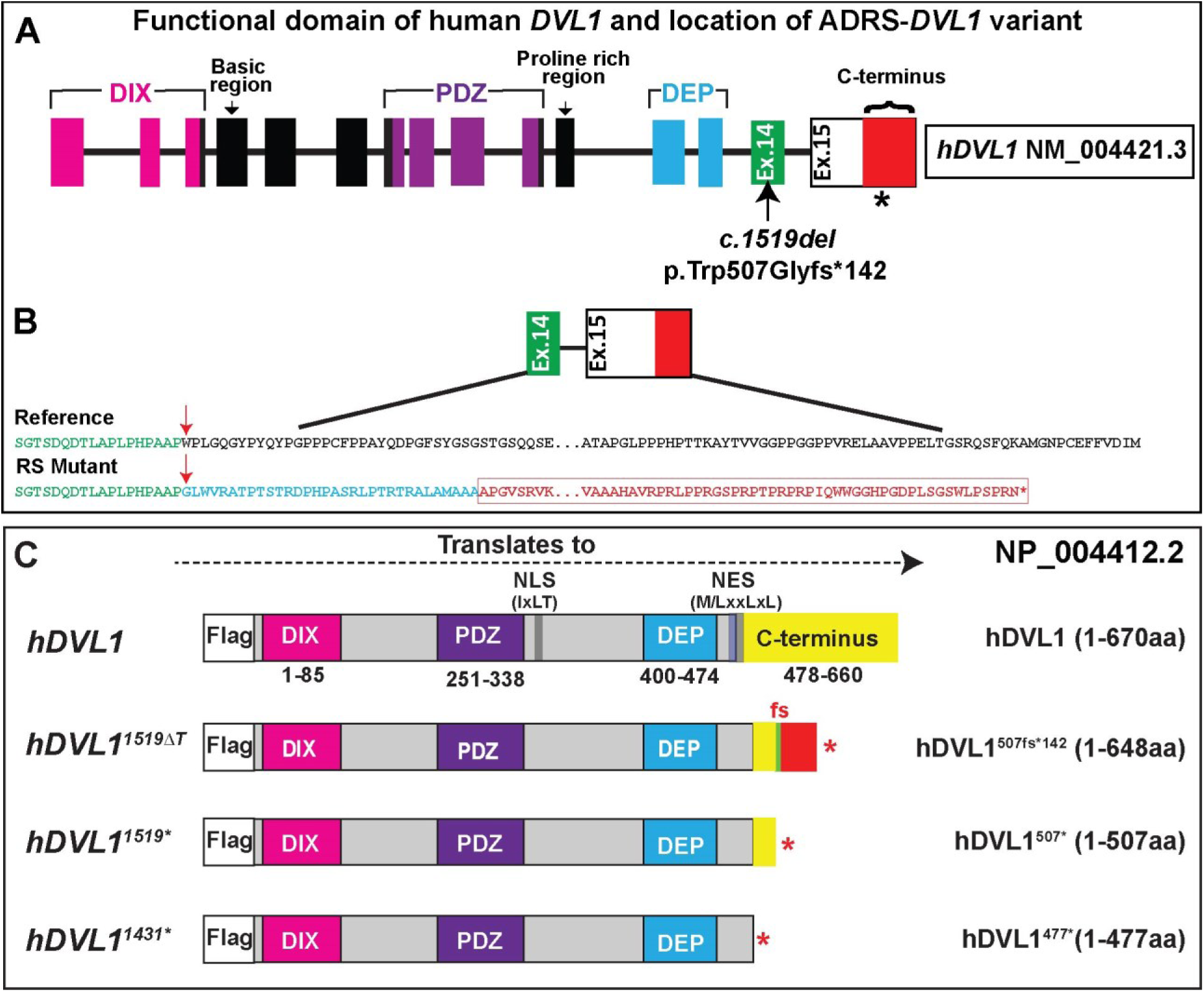
Illustration of hDVL1 constructs and western blot analysis. (A) The positions of functional domains (DIX, PDZ, DEP), nuclear localization sequence (NLS, IxLT), nuclear export sequence (NES, M/LxxLxL), and the C-terminus (yellow) are indicated. The six amino acid NES is located in the DEP and the C-terminus (blue). The h*DVL1*^1519ΔT^ (hDVL1^507fs*142^) construct displays the frameshift (green line), abnormal C-terminus (red), and location of the STOP codon. In h*DVL1*^1519*^(hDVL1^507*^), the STOP codon was introduced after the frameshift. Lastly, the h*DVL1*^1431*^ (hDVL1^477*^) has a STOP codon inserted three amino acids after the DEP-domain, has lost the entire C-terminus. A FLAG-tag (DYKDDDDK) was added at the N-terminus for all constructs.

The *DVL1* contains 15 exons (Fig. 1A) and is on human chromosome 1p36.3. The 12 identified Robinow syndrome *DVL1* variants cluster within the penultimate and the last exon (exon 14 and 15), generating frameshifts that result in an abnormal C-terminus (Fig. 1A,B, Table S3). Depending on the position of the frameshift mutation, at least 109 novel amino acids replace at least 68 amino acids of the normal C-terminus (Fig. 1A,B)(Hu et al., 2022; Lima et al., 2022; White et al., 2015; Zhang et al., 2022). Interestingly, the novel C-terminal peptide is common to all identified ADRS-*DVL1* individuals (Hu et al., 2022; White et al., 2015; Zhang et al., 2022) (Fig. 1A-C). The last 23 amino acids of DVL1 C-terminus are evolutionarily conserved (Mlodzik, 2016; Qi et al., 2017). The extreme end of the DVL C-terminus loops back to bind to its own PDZ domain thus maintaining the ‘closed’ conformation of DVL leading to autoinhibition and reduced activity of the WNT pathway (Bunn et al., 2015; Qi et al., 2017). The non-canonical WNT pathway was also moderately reduced in *Xenopus* embryos when the DVL1 protein formed a closed conformation (Qi et al., 2017). The C-terminus has been shown to antagonize canonical signaling stimulated by β-catenin in a *Ror2*-dependent manner (Witte et al., 2010). Less is known about the C-terminus since many structure-function studies do not specifically delete this domain (Rothbacher et al., 2000; Sokol, 1996; Tauriello et al., 2010; Tauriello et al., 2012). The effects of the novel C-terminal peptide on DVL1 subcellular localization, protein conformation, protein-protein interaction, WNT signal transduction and transcription are unknown.

Here we focused on the craniofacial phenotypes, one of the most penetrant aspects of ADRS, whether caused by *WNT5A, FZD2* or *DVL1* mutations (Zhang et al., 2022). To understand how the ADRS h*DVL1^1519^*^Δ*T*^ variant differs from the wild-type (wt) h*DVL1*, our strategy was to overexpress the wt gene and compare the phenotypes to embryos expressing h*DVL1^1519^*^Δ*T*^ (Gignac et al., 2019; Gignac et al., 2023; Hosseini-Farahabadi et al., 2017). Through expression of the human gene on top of the chicken genome, we mimic the heterozygous, ADRS genotype within the chicken embryo model. The pathogenicity of h*DVL1* variants was previously tested in developing limbs of chicken embryos and in *drosophila* wings (Gignac et al., 2023). The h*DVL1^1519^*^Δ*T*^ (p.Trp507Glyfs*142) was found to produce highly penetrant limb and wing phenotypes (Gignac et al., 2023). h*DVL1^1519^*^Δ*T*^ produces a stable mRNA product that translates into a mutant protein (Bunn et al., 2015). We therefore focused on investigating the effects of h*DVL1^1519^*^Δ*T*^ on developing face in chicken embryos. To characterize the effects of the abnormal C-terminal peptide produced by h*DVL1^1519^*^Δ*T*^ (hDVL1^507*fs^), we designed two additional constructs. In the first construct, a STOP codon (TAA) was inserted after nucleotide 1519 (h*DVL1^1519*^*, hDVL1^507*^)(Bunn et al., 2015). In the second construct, the STOP codon was inserted further upstream after nucleotide 1431, and lacks the entire C-terminus (h*DVL1^1431*^*; hDVL1^477*^). The validation of the translation of all h*DVL1* constructs into stable proteins of the anticipated size was conducted with Western Blots and quantified in ImageJ (Fig. S1)(Stael et al., 2022).

We assessed the morphogenetic, skeletogenic and biochemical impacts triggered by the *GFP* (control) and four h*DVL1* (wt h*DVL1*, h*DVL1^1519^*^Δ*T*^, h*DVL1^1519*^*, h*DVL1^1431*^*, N-terminal FLAG-tag) constructs, aiming to elucidate the significance of the novel C-terminus associated with ADRS and to gain deeper insights into the general role of the h*DVL1* C-terminus. We have successfully recapitulated the facial widening and bone hypoplasia phenotypes caused by h*DVL1^1519^*^Δ*T*^ variant. In addition, we have shown that the variant is weak in activating the WNT canonical and non-canonical pathways. This is the first study to establish a link between the abnormal C-terminal sequence in *DVL1* variant with RS phenotypes. Furthermore, out identified that the first 30 amino acids of the C-terminus are crucial for development.

## Results

### Robinow syndrome associated h*DVL1* variant inhibits upper beak patterning

To access the upper beak phenotypes, embryos were injected in the frontonasal mass at stage 15 [Embryonic day (E) 2.5] with *GFP*, wt or h*DVL1 ^1519^*^Δ*T*^ viruses and incubated until full skeletal maturation (stage 38, E12.5, Table S5). As anticipated, the skulls of embryos infected with the *GFP* virus, exhibited a normal size upper beak capped with an egg tooth (Fig. 2A). Embryos injected with wt h*DVL1* and h*DVL1 ^1519^*^Δ*T*^ viruses displayed a shortened and deviated upper beak and the egg tooth was present (Fig. 2B, C). Analysis of alcian blue (to detect cartilage) and Alizarin red (to detect mineralized bone) stained skeletal preparations showed that *GFP* embryo has normal shape, size and ossification of frontonasal mass derived bones (Fig. 2**Error! Reference source not found.**A-A’’’). In contrast embryos injected with wt *DVL1* (Fig. 2B’-B’’’; Fig. S2A-C’) or h*DVL1 ^1519^*^Δ*T*^ viruses (Fig. 2C’-C’’’; Fig. S2D-F’) produced curved, shorter upper beaks, particularly affecting the premaxillary bone. Moreover, analysis from the palatal side revealed that the premaxillary bone, originating from the frontonasal mass was absent or unossified, specifically on the injected side (Fig. 2B’’’, C’’’). This observation was noted in embryos injected with wt *DVL1* (57%) or *DVL1 ^1519^*^Δ*T*^ (92%) viruses (Table S5, Fig. 2 B’’’, C’’’). Notably, there was no difference in alizarin red staining between the viruses suggesting that the ossification of the upper beak bones was not affected. Our data implies that overexpression of h*DVL1* during the development of the chicken embryos face can result in patterning defects. Intriguingly, the penetrance of the beak phenotype was significantly higher in embryos infected with the *DVL1 ^1519^*^Δ*T*^ (Fig. 2D,E,F, Fig. S2D-F’). The observed patterning defects may contribute to the midface hypoplasia observed in ADRS.

**Figure 2.**
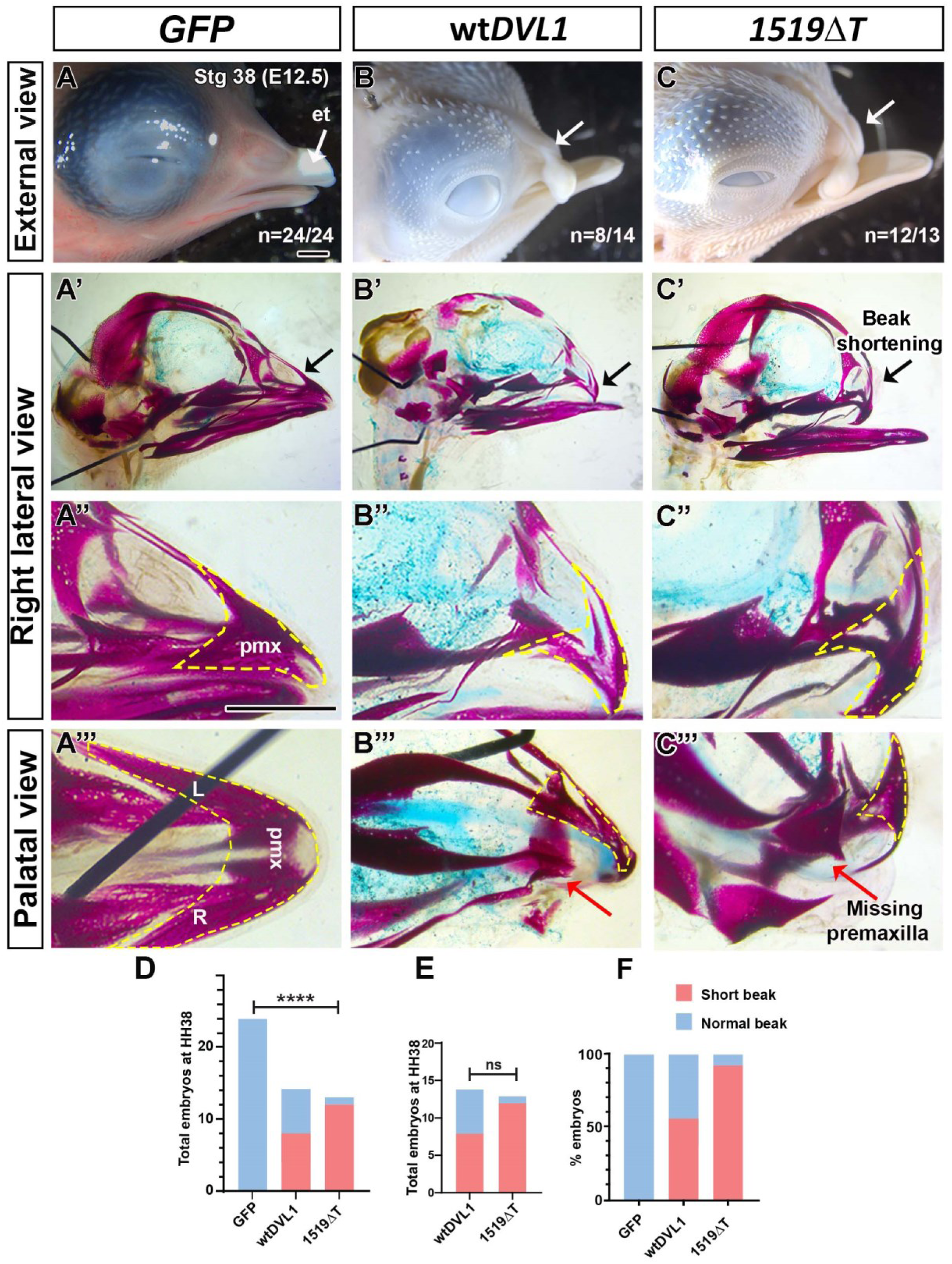
Upper beak phenotypes induced after overexpression of RCAS:GFP, RCAS: wt hDVL1 and RCAS:hDVL1^1519ΔT^. Embryos injected with RCAS:GFP, wt hDVL1, DVL1^1519ΔT^ variant into the right frontonasal prominence at stage 15 (E2.5) and fixed 10 days post injection at stage 38 (E12.5). (A) External photographs of stage 38 embryos in right lateral view revealed normal upper beak morphology in those injected with the GFP virus, whereas (B, C) embryos injected with wt or mutant hDVL1 viruses exhibited relative shortening and deviation of the upper beak, indicated by white arrows. (A’) Wholemount skulls stained with alcian blue and alizarin red showed a normal upper beak skeleton in GFP-injected specimens (24/24). (B’, C’) In contrast, embryos injected with wt (n=8/14) or mutant hDVL1 (n=12/13) viruses displayed hypoplastic and abnormal morphology of upper beak bones, as indicated by black arrows. (A’’) High-power views of the premaxillary bone (marked by yellow dash line) demonstrated normal patterning in GFP injected embryos and (B’’. C’’) abnormal patterning and inhibition of ossification in the upper beak due to the overexpression of hDVL1 viruses. (A’’’) Palatal view of the upper beak showing GFP treated embryos showed a normal premaxillary bone while (B’’’, C’’’) embryos treated with hDVL1 viruses exhibited a missing right premaxillary bone, indicated by the black arrow. (D) Contingency analysis followed by Fisher’s exact test and (E) Fraction of total analysis showing proportion of embryos with beak phenotype performed with GraphPad Prism 10.1.0. The faint cartilage stain (blue) in all skulls is a technical error. Key: pmx – premaxilla. Scale bar: A-C’, A’’-C’’’ = 2mm.

### ADRS variant impedes narrowing of the frontonasal mass and differentiation of the prenasal cartilage

The early stages of facial morphogenesis are conserved between human and chicken embryos (Abramyan and Richman, 2018). A subset of embryos was collected immediately after the onset of chondrogenesis for histological analysis and immunostaining (stage 29, E6.5). Since ADRS patients present with wide face, we measured the width of the frontonasal mass (Illustration in Fig. 3A) using histological sections (Fig. 3D-F). Interestingly, the embryos injected with h*DVL1 ^1519^*^Δ*T*^ had significantly wider frontonasal mass compared to wt h*DVL1* and *GFP* (Fig. 3A). In addition, *GFP* (n=5) and wt h*DVL1* (n=4) injected specimens showed normal morphology of the prenasal cartilage while h*DVL1^1519^*^Δ*T*^ (n=8) caused unilateral inhibition of the prenasal cartilage (Fig. 2D-F). Interestingly, in the *GFP* and wt h*DVL1* embryos presence of virus (antibody detects viral GAG protein) did not affect the prenasal cartilage (Fig. 2D,D’,E,E’). In contrast, h*DVL1^1519^*^Δ*T*^ virus caused inhibition of the prenasal cartilage (Fig. 2F, F’). Next, we probed serial sections with SOX9 (marker for early cartilage) and TWIST1 (marks undifferentiated mesenchyme). The *GFP* or wt h*DVL1* viruses had no effect on SOX9 and TWIST1 expression (Fig. 2G,H). In h*DVL1 ^1519^*^Δ*T*^ embryos, the SOX9-negative mesenchymal cells in the prenasal cartilage were islands of undifferentiated TWIST1-positive mesenchymal cells(Fig. 2I). This suggests that ADRS *DVL1^1519^*^Δ*T*^ variant stalls the frontonasal mass mesenchyme in an undifferentiated state, inhibits skeletogenesis leading to loss of the premaxillary bone and beak deviation (Fig. 2C-C’’’). Next, we hypothesized that the truncation of the beak could be due to decreased cell proliferation on the injected side of the frontonasal mass. The h*DVL1^1519^*^Δ*T*^ virus significantly inhibited cell proliferation compared wt h*DVL1* (Fig. 2B,J-LB). Since *DVL1* functions in canonical pathway, we quantified expression of WNT canonical pathway marker, nuclear β-catenin. There was no differences between the viruses (Fig. 2C,M-O). Overall, the immunostaining data suggests that inhibition of chondrogenesis and cell proliferation may be contributing to the short and deviated upper beak in h*DVL1^1519^*^Δ*T*^.

**Figure 3.**
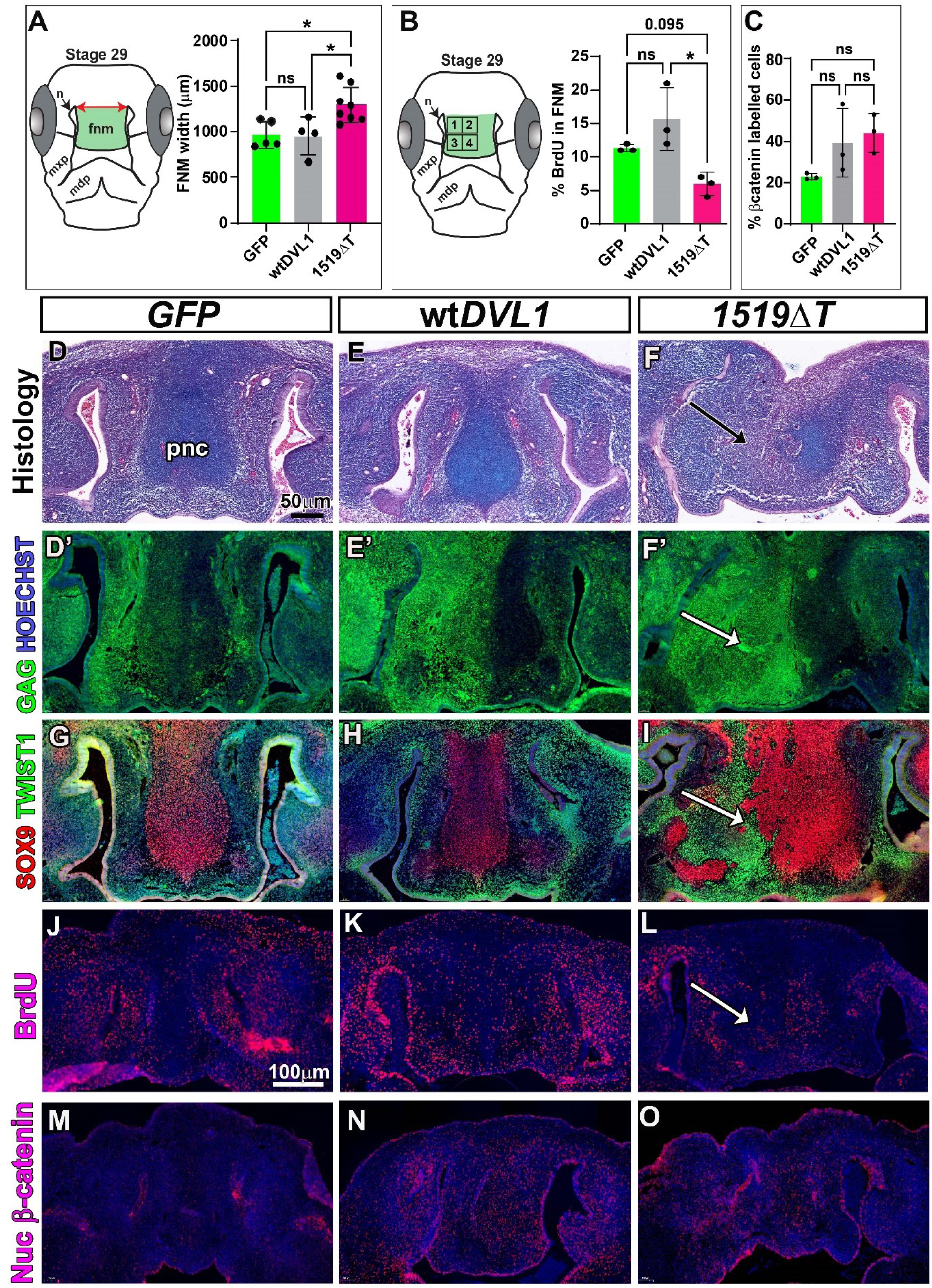
Analysis of embryos injected with GFP, wt hDVL1 and hDVL1^1519ΔT^ at stage 28.5 (96h post-injection, E6.0) Embryos injected into the right frontonasal mass with RCAS viruses containing GFP or hDVL1 were fixed in 4% PFA 96h post-injection and sectioned frontally. (A) Frontonasal mass width measurements and statistical analysis (1-way ANOVA, Tukey’s). (B) BrdU counting scheme and percentage labeled cells for all 4 regions combined (1-way ANOVA, Dunnett’s). (C) Proportion of frontonasal mass mesenchyme expressing nuclear β-catenin showed significantly inhibited proliferation (one-way ANOVA). (D-F) Sections of the head were stained with Alcian blue and Picrosirius red. (D,E) Normal prenasal cartilage in GFP and wt hDVL1 infected embryos (F) while the cartilage is abnormal and absent on the right side in the embryos injected with hDVL1^1519ΔT^. (D’-F’) Adjacent sections were stained with anti-GAG antibody and counterstained with Hoechst to locate the virus in the tissues. (G-I) Near-adjacent sections stained with SOX9 (stains chondrocyte nuclei) and TWIST (stains undifferentiated mesenchymal cell nuclei) antibodies. (G,H) In GFP and wt hDVL1 infected embryos SOX9 was expressed in the prenasal cartilage. (I) the hDVL1^1519ΔT^ variant led to TWIST1 expression displacing areas that normally are SOX9 positive. (I-K) Proportion of BrdU-positive cells in the virus infected area showed low proliferation in differentiating cartilage in hDVL1^1519ΔT^ virus. (L) Immunostaining for nuclear βcatenin was performed on a different set of GAG-positive embryos. Normally there is very little expression of nuclear βcatenin in the cartilage as observed in embryos injected with GFP. (M, N) There appeared to be increased expression in wt hDVL1 and variant DVL1 although the changes were not statistically significant (quantified, data not shown). Key: FNM – frontonasal mass, n – nasal slit, ns – non-significant, mdp – mandibular prominence, mxp – maxillary prominence, pnc – prenasal cartilage. P values: *= p<0.05, ns – not significant. Scale bar in C-H = 50µm, same scale bar applies to I-N = 100µm.

To further explore the effects of *DVL1* variant, we measured gene expression changes 48 h post-injection (Fig. S3A-E). The levels of h*DVL1* were 3-to-6-fold higher in embryos injected with h*DVL1* viruses while the g*DVL1* remained unchanged (Fig. S3A,B). The g*CTNNB1* was significantly downregulated by h*DVL1^1519*^* (Fig. S3B) while h*DVL1^1519^*^Δ*T*^ variant decreased levels of SOX9 compared to *GFP* (Fig. S3D). The RNA levels across other WNT and skeletogenic markers genes were not significantly different than *GFP* (Fig. S3B, D) controls or wt h*DVL1*(Fig. S3C, E). Although there were no significant alterations in other genes, we cannot rule out the possibility that significant alterations may be seen at later stage of development.

### Abnormal C-terminus in mutant h*DVL1* contributes to observed phenotypes

Next, we investigated the association between the observed phenotypic changes and the novel C-terminal peptide with virus containing *DVL1^1519*^*or *DVL1^1431*^*. The interpretation of observed effects will be based on the following criteria; (1) Similarities between the wt h*DVL1* and the h*DVL1^1519*^*or h*DVL1^1431*^* would imply that the observed phenotypes are caused due to overexpression of h*DVL1*, (2) if h*DVL1^1519^*^Δ*T*^ resembles h*DVL1^1519*^* it would suggest that the phenotypes are not associated with the presence of abnormal C-terminus, (3) distinct effects caused exclusively by the h*DVL1^1519*^* or by h*DVL1^1431*^* could indicate neomorphic effects, facilitating the clarification of the effects of the novel C-terminal peptide. Surprisingly, embryos injected with h*DVL1^1519*^*exhibited normal beak development (Fig. 4A-A’’). In contrast, h*DVL1^1519^*^Δ*T*^, with abnormal C-terminal sequence, had a major impact on beak patterning (Fig. 2C-C’’’; **Error! Reference source not found.**D-F’). This suggests that the h*DVL1^1519*^*protein lacking a part of the C-terminus allows normal beak formation while addition of the novel amino acids in the ADRS variant (*DVL1^1519^*^Δ*T*^) likely causes a gain-of-function (Fig. 4A-A’’).

**Figure 4.**
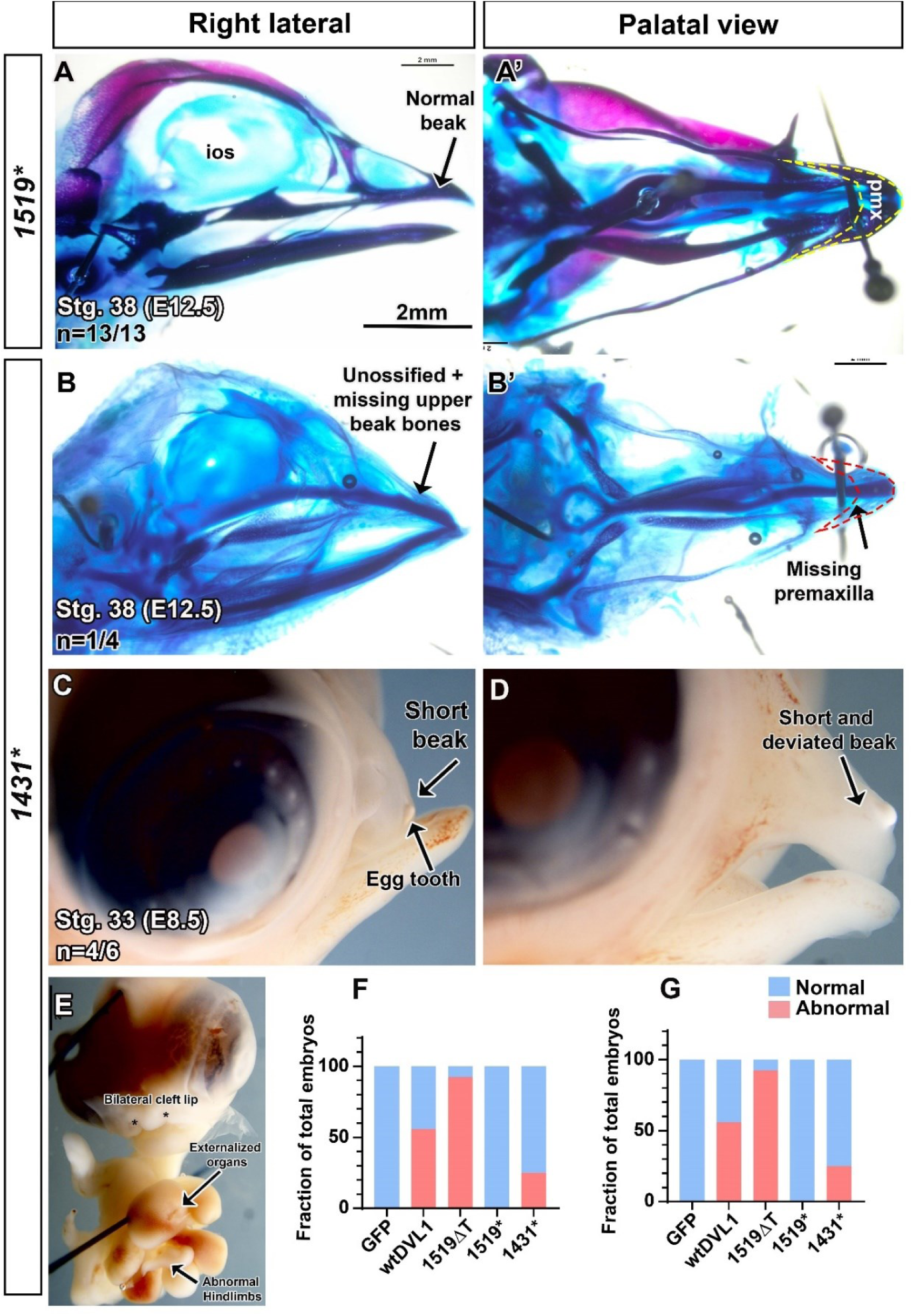
Wholemount specimen treated with RCAS:hDVL1^1519*^ and RCAS:hDVL1^1431*^. (A) Wholemount skull injected with DVL1^1519*^ (n=13/13) stained with alcian blue (stains cartilage) and alizarin red (stains bone) shows normal upper beak morphology. (A’) Palatal view showing normal premaxillary bone (yellow dash line) was ossified and had normal morphology. (A) Wholemount skull injected with DVL1^1431*^ (n=1/4) stained with alcian blue (stains cartilage) and alizarin red (stains bone) shows completely unossified upper beak at stage 38. None of the skeletal elements are clearly visible except for the prenasal cartilage, nasal bone is missing. (A’) Palatal view showing that the premaxillary bone is missing. (B, C) Individual embryos collected at stage 33 (E8.5) show that the upper beak was short and deviated in 4/6 embryos. (D) Embryos that did not survive still 6 days were collected for phenotypic analysis show bilateral cleft lip, hindlimb defects and externalized organs. (E) Contingency analysis followed by Fisher’s exact test Fraction of total analysis (bottom graph) showing proportion of embryos with beak phenotype performed with GraphPad Prism 10.1.0. Scale bar: A-D = 2mm

The next step was to evaluate the effects of *DVL1* C-terminus (h*DVL1^1431*^* virus) in beak development and comparing these phenotypes with those observed in connection with in vivo overexpression of other h*DVL1* viruses. However, unlike all the other viruses studied to date in the Richman lab (Ashique et al., 2002; Geetha-Loganathan et al., 2014; Gignac et al., 2019; Gignac et al., 2023; Hosseini-Farahabadi et al., 2013; Hosseini-Farahabadi et al., 2017; Nimmagadda et al., 2015), a significant majority of embryos injected with the h*DVL1^1431*^*virus died by stage 38 (day 10) (TableS5). Among the limited number of embryos that survived to stage 38, one out of four embryos showed an unossified skull and the absence of upper beak bones (Fig. 4B, B’). For phenotypic analysis, embryos were collected at stage 33 (E8.5). The survival rate was 42% (TableS5). Four out of six surviving embryos at stage 33 displayed a short upper beak, resembling the phenotype seen with h*DVL1^1519^*^Δ*T*^ (Fig. 4C,D). Examination of embryos that did not survive up to stage 33 revealed defects in face and other organs (Fig. 4E). The hindlimb development appeared stunted and the heart was located outside the body due to spread of virus to adjacent areas like the brain, heart, or neural tube, located close to the frontonasal mass (Fig. 4E). This spread might have led to inhibition of somite development and the caudal extension of the embryo. Taken together, our in vivo overexpression of h*DVL1* lacking the C-terminus (h*DVL1^1431*^*) can disrupt development in chicken embryos.

### *DVL1^1519^*^Δ*T*^ and *DVL1^1431*^*inhibit chondrogenesis in frontonasal mass micromass cultures

Since the ADRS variant h*DVL1^1519^*^Δ*T*^ inhibited chondrogenesis in vivo, we investigated exactly which stage of chondrogenesis was affected using frontonasal mass micromass cultures. On day 4, there were no noticeable differences observed in cartilage sheets between the different viruses showing the early stages of cartilage condensation were not affected (Fig. 5A-E). However, by day 6, viruses *DVL1^1519^*^Δ*T*^ and *DVL1^1431*^* induced qualitative reductions in chondrogenesis compared to control virus *GFP*, wt *DVL1* (Fig. 5F-J). A further decrease in cartilage was visible in day 8 cultures infected with *DVL1^1519^*^Δ*T*^ and *DVL1^1431*^* but not for *DVL1^1519^* which was not changed compared to controls (Fig. 5K-O).

**Figure 5.**
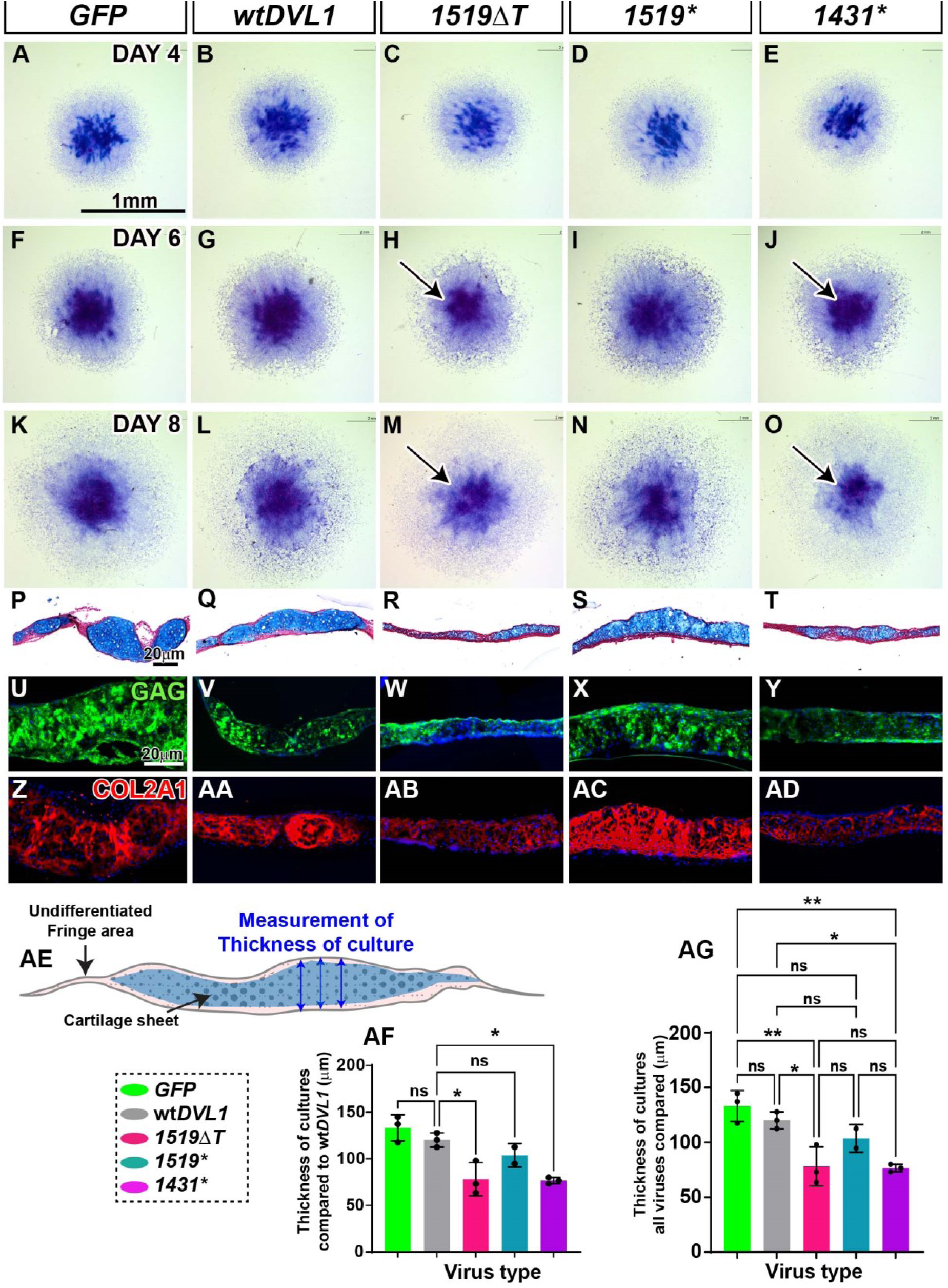
Effects of h*DVL1* viruses on frontonasal mass micromass cultures. Mesenchymal cells harvested from the frontonasal mass of stage 24 (E4.5) embryos were plated into high density (2 x 107 cells/mL) cultures. Cultures fixed on (A-E) day 4, (F-J) day 6 and (K-O) day 8 stained in wholemount with alcian blue (stains cartilage) and counterstained with hematoxylin. (A-O) Cartilage sheets derived from frontonasal mass mesenchyme can be seen in the center of the culture. (H, J, M, O) Note the quantitative reduction in chondrogenesis (blue stain) in cultures infected with *DVL1^1519^*^Δ*T*^ and *DVL1^1431*^* on day 6 and day 8. (P-T) A subset of day 8 cultures were fixed in 4% PFA, embedded in paraffin and sectioned for histology and immunostaining. (AE - AG) Histological sections stained with alcian blue and picrosirius red and the thickness of cultures (n=3) was measured in CaseViewer. (P-T, AF, AG) *DVL1^1519^*^Δ*T*^ and *DVL1^1431*^* virus treated cultures were significantly thinner compared to GFP, wt *DVL1*, and *DVL1^1519*^*. Statistical analysis done with one-way ANOVA followed by (AF) Dunnett’s multiple comparison test or (AG) Tukey’s post-hoc test. Scale bar in A to O = 100µm, P-T= 20µm. P values *= p<0.05, **= p< 0.01, ns-not significant

In order to delve deeper into the impact of the viruses, a subset of day 8 cultures was fixed for subsequent histological analysis. Histological analysis revealed variations in thickness among cultures, particularly the much thinner cultures infected with *DVL1^1519^*^Δ*T*^ and *DVL1^1431*^* (Fig. 5P-T). Further measurements conducted on the thickest areas within these cultures (Fig. 5AE) confirmed a significant reduction in thickness for those infected with *DVL1^1519^*^Δ*T*^ and *DVL1^1431*^* in comparison with *GFP* and wt *DVL1* (Fig. 5AF). Interestingly, *DVL1^1519*^* viruses had similar thickness to control cultures (Fig. 5AF). All the cultures showed abundant spread of virus (Fig. 5U-Y). Next, we examined how the viruses affected expression of the most abundant protein found in cartilage, type II collagen (COL2A1). All cultures exhibited a typical expression pattern of COL2A1, mirroring the observations seen with alcian blue staining in the histological sections (Fig. 5Z-AD). Collectively, the findings from the *in vitro* experiments mirrored the in vivo results, demonstrating that the *DVL1^1519^*^Δ*T*^ and *DVL1^1431*^* viruses hindered chondrogenesis but that the *DVL1^1519*^* virus allowed development to proceed normally.

### *DVL1^1519^*^Δ*T*^ and *DVL1^1431*^*cause downregulation of chondrogenic markers in frontonasal mass cultures

In RNA expression analysis in the micromass cultures, we confirmed that human *DVL1* levels were 10-to-15-fold higher than *GFP* (Fig. S5A) however there was no effect on *Gallus DVL1* or evidence of a feedback loop (Fig. S5B). Cultures of h*DVL1^1519^*^Δ*T*^ and h*DVL1^1431*^* exhibited diminished RNA levels of early chondrogenic markers, specifically *gCOL2A1* and *gSOX9,* in contrast to *GFP cultures* (Fig. S5B). The expression level of g*TWIST2,* a mesenchymal cell differentiation marker, was significantly lower in cultures infected with h*DVL1^1519^*^Δ*T*^ and h*DVL1^1431*^* compared to those with wt *DVL1* and *GFP* cultures (Fig. S5B, C). In all these analyses, the h*DVL1^1519*^* had expression that was not significantly different than wt*DVL1*. Since the expression of *COL2A1* was lower in cultures infected with h*DVL1^1519^*^Δ*T*^ and h*DVL1^1431*^*, we measured expression of g*MMP13* considering its role in degradation of cartilage matrix (Hu and Ecker, 2021). A previous study from our lab showed that treatment of mandibular micromass cultures to *WNT5A* conditioned media caused inhibition of chondrogenesis due to increased g*MMP13* expression (Hosseini-Farahabadi et al., 2013).

Interestingly, the g*MMP13* transcript was significantly downregulated by h*DVL1^1519^*^Δ*T*^ and h*DVL1^1431*^*compared to wt h*DVL1* (Fig. S5C). This suggests that the mechanism leading to reduced cartilage in h*DVL1^1519^*^Δ*T*^ and h*DVL1^1431*^* infected cultures is not due to a degradation of cartilage matrix. Instead, reduction in cartilage may be due to failure of the chondrocytes to secrete sufficient cartilage matrix. The reduced g*MMP13* levels can be a response of the cells to the reduced cartilage matrix in the h*DVL1^1519^*^Δ*T*^ and h*DVL1^1431*^*, but is not a mechanism directly contributing to the observed phenotype.

Since h*DVL1* functions in the WNT canonical pathway which regulates chondrogenesis, we analyzed WNT pathway genes including g*CTNNB1*, gA*XIN2*, g*LEF1*, g*FZD2*, g*WNT5A* and g*ROR2*. The level of g*CTNNB1* was significantly upregulated by h*DVL1^1519*^*compared to *GFP* but remained comparable to other h*DVL1* viruses (Fig. S5B). g*LEF1* and g*FZD2* were downregulated by h*DVL1^1519^*^Δ*T*^ compared to wt h*DVL1* (Fig. S5C). The levels of g*LEF1* and g*FZD2 DVL1^1431*^* showed similar trend like *DVL1^1519^*^Δ*T*^ but were not significantly different compared to the wt h*DVL1* (Fig. S5C). This could explain the similarity in the phenotypes induced by *DVL1^1431*^*and *DVL1^1519^*^Δ*T*^. Since the bone morphology was severely affected, we investigated whether the mutant *DVL1* viruses initiates crosstalk with the BMP pathway. We specifically measured the effects on g*BMP2* which promotes chondrogenesis in the micromass culture system (Fischer et al., 2002) and g*MSX1,* a homeobox gene expressed abundantly during craniofacial development (Alappat et al., 2003). Interestingly, only h*DVL1^1519*^*upregulated g*BMP2* and g*MSX1* compared to wt h*DVL1* (Fig. S5D). The differences between the RT-PCR data in vivo (Fig. S3) and *in vitro* (Fig. S5) may be attributed to the higher viral load at the time of infection in vitro compared to in vivo. The micromass cultures exclude epithelium which may change the responses of cells to the viruses. Additionally, the timing of molecular changes in vivo may have occurred later than in vitro. Thus far, shortening and deviation of the upper beak, inhibition of chondrogenesis and RNA expression profile in micromass cultures have demonstrated that the effects induced by the h*DVL1^1519^*^Δ*T*^ are similar to h*DVL1^1431*^*.

### *DVL1* C-terminus is crucial for both branches of WNT signaling pathway

DVL proteins play a vital role in the canonical WNT pathway signaling (Gao and Chen, 2010), therefore we utilized SuperTopflash (STF) luciferase assays with plasmids that contain the response element for β-Catenin signaling to measure canonical activity. We used the ATF2 plasmid to measure JNK-PCP signaling (Geetha-Loganathan et al., 2014; Ohkawara and Niehrs, 2011). In frontonasal mass primary cell cultures wt h*DVL1* strongly activated STF activity, while the h*DVL1^1519^*^Δ*T*^ variant exhibited weaker activation (Fig. 6A). Similarly, h*DVL1^1519^*^Δ*T*^ demonstrated significantly weak ATF2 activation compared to wt h*DVL1* (Fig. 6B). This suggests that the abnormal C-terminal peptide caused by the frameshift leads to a conformational change that possibly affects the interaction of the C-terminus and the PDZ domain thus affecting the WNT signaling (Itoh et al., 2005; Qi et al., 2017). Additionally, this data confirms that the luciferase activity of the h*DVL1^1519^*^Δ*T*^ is similar in chicken frontonasal mass mesenchymal cells and HEK293T cells, as published (Gignac et al., 2023). Both h*DVL1^1519*^* and h*DVL1^1431*^*were weaker in activating STF and ATF2, similar to h*DVL1^1519^*^Δ*T*^. This suggests that the C-terminus sequence after nucleotide 1519 is crucial in regulating both canonical and non-canonical WNT signaling pathways

**Figure 6.**
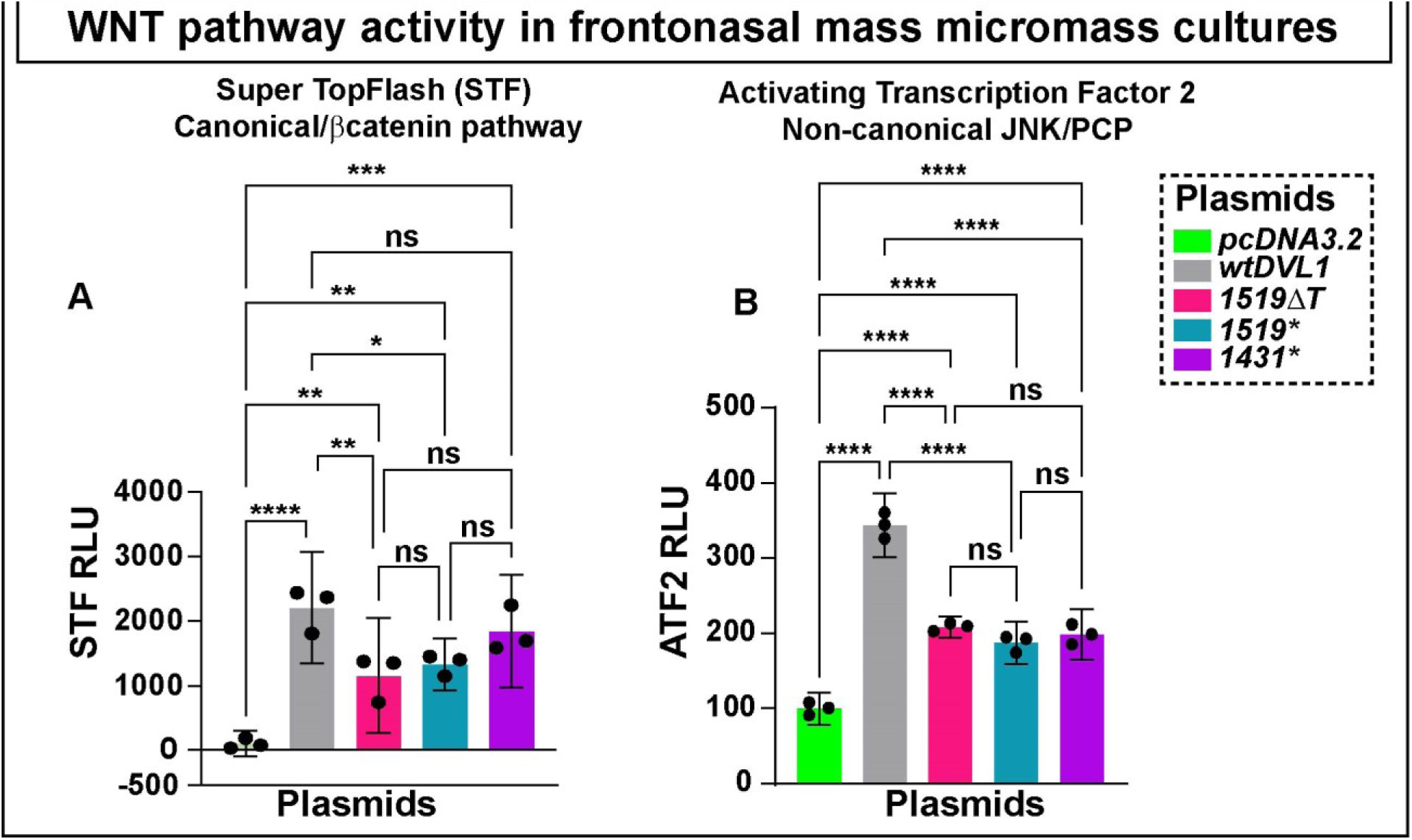
Relative luciferase reporter activity in micromass cultures following transient transfection of pcDNA3.2 or h*DVL1* plasmids. The activation of (A) STF and (B) ATF2 reporters was significantly lower in *DVL1^1519^*^Δ*T*^, *DVL1^1519*^* and *DVL1^1431*^* compared to wt h*DVL1*. Three biological replicates, three technical replicates and the experiment was repeated three times. Statistics done with one-way ANOVA followed by Tukey’s post hoc test multiple comparison test using Prism 10.1.0. P values *= p<0.05, **= p< 0.01, ***=p<0.001, ****=p<0.0001, ns-not significant.

### h*DVL1*^1519**Δ**T^ protein accumulates within the nucleus in HEK293T DVL-TKO cells

The C-terminus of DVL contains a nuclear localization sequence (NLS) and a nuclear export sequence (NES, Fig. 1C) that gives DVL proteins the ability to shuttle between the cytoplasm and the nucleus (Itoh et al., 2005; Pruller et al., 2022). The leucine-rich NES in Dsh was shown to be located between 510-515 (Itoh et al., 2005) but the exact location in hDVL1 is not known. Based on the sequence (M/LxxLxL), the NES in hDVL1 occupies a part of DEP domain and a part of the C-terminus of the DVL protein (Fig. 1C). The shuttling of DVL proteins is crucial for WNT signaling (Itoh et al., 2005; Weitzman, 2005). The movement of DVL proteins into the nucleus is controlled by FOXK transcription factors (Wang et al., 2015). Additionally, the C-terminal domain of DVL proteins has been shown to be essential for protein-protein interactions (Gan et al., 2008; Gloy et al., 2002; Goto et al., 2013; Harnos et al., 2019; Hikasa and Sokol, 2004). Our objective was to assess whether the presence (wt hDVL1, hDVL1^1519*^) or absence (hDVL1^1519ΔT^, h*DVL1^1431*^*) of NES in the hDVL1 constructs impacts the subcellular protein localization. Here, we used HEK293T cells lacking all three *DVL* isoforms (DVL Triple Knockout; referred to as HEK293T DVL-TKO). In these cells all three human *DVL* paralogs were deleted by CRISPR–Cas9 technology (Cervenka et al., 2016). Due to the unavailability of effective DVL1 antibodies, the use of Flag tags allowed us to monitor hDVL1 localization in fixed cells 48 hours after transfection in the HEK293T DVL-TKO cells.

The wt hDVL1 and hDVL1^1519*^ were well distributed in the nucleus, cytoplasm or both, and formed distinct cytoplasmic puncta (Fig. 7A,C,E,F). Thus, the wt DVL1 and the 1519* protein showed similar subcellular localization despite of the absence of the most terminal part of the C-terminus. Interestingly, the hDVL1^1519ΔT^ protein was primarily localized within the nucleus (Fig. 7B, E, F). As expected, cells transfected with h*DVL1^1431*^* (lack the NES) had 83% cells with hDVL1 protein in the nucleus (Fig. 7D). Both wt hDVL1 and hDVL1^1519*^ showed clearly identifiable cytoplasmic puncta which were almost completely absent in hDVL1^1519ΔT^ (Fig. 7E). This suggests that the ADRS hDVL1^1519ΔT^ protein has the tendency for nuclear retention, possibly due to the loss or interference of the abnormal C-terminus with the NES or other structural changes within the hDVL1 protein. The effects of the mutation on the structure of DVL1 remains undetermined.

**Figure 7.**
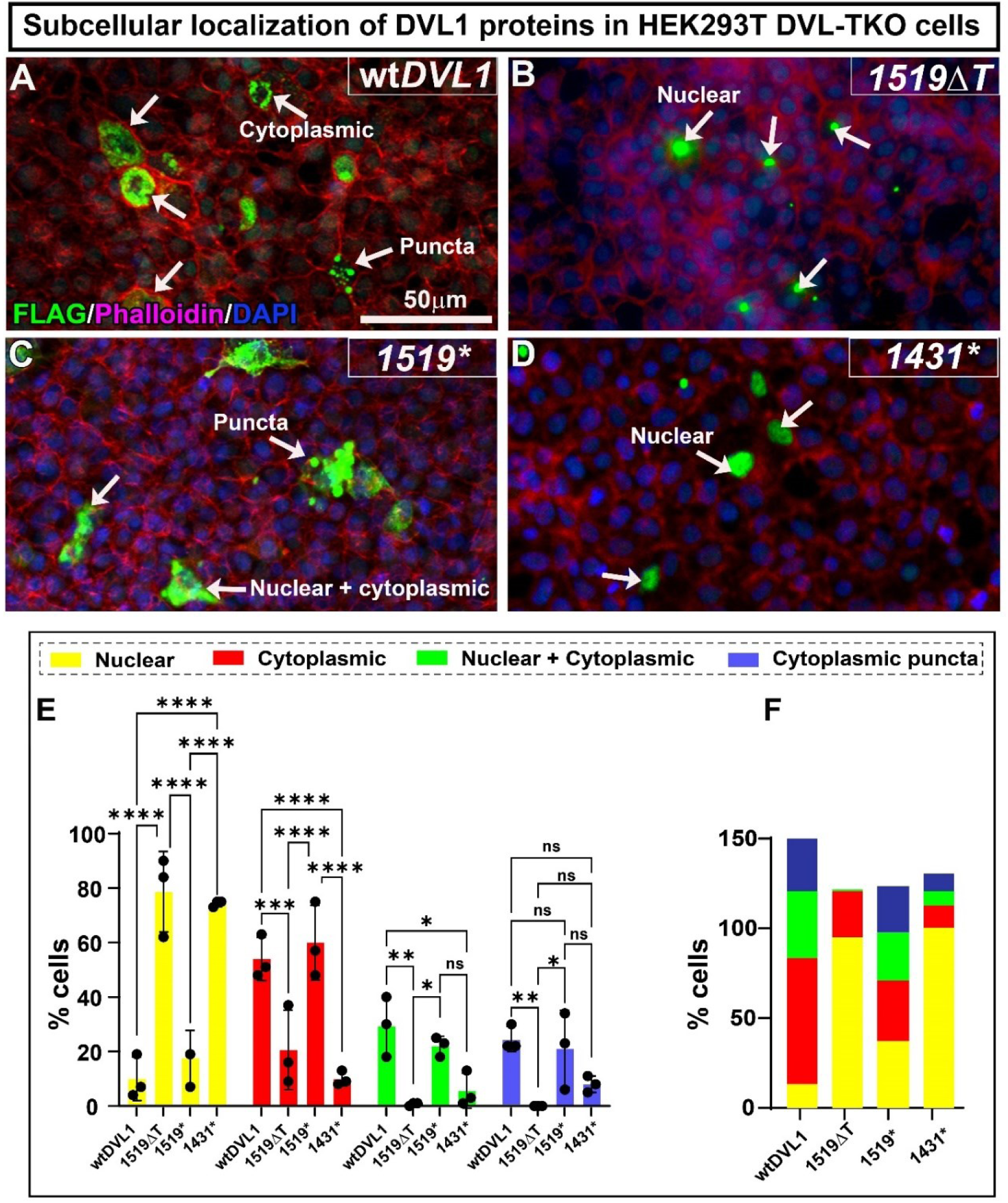
Expression of FLAG-hDVL1 protein in HEK293T DVL-TKO cells. Cells were transfected using lipofectamine combined with Flag-tagged hDVL1 constructs, fixed 48h post-transfection and stained with anti-Flag and Phalloidin. Cells expressing FLAG are highlighted with white arrows. (A) The distribution of Flag-tag wt hDVL1 protein appeared throughout the nucleus and cytoplasm with a distinctive punctate appearance. (B) The ADRS-hDVL1^1519ΔT^ protein exhibited a higher proportion of transfected cells with FLAG localized within the nucleus. (C) In contrast, the hDVL1^1519*^ protein displayed subcellular localization similar to wt hDVL1. (D) The hDVL1^1431*^ construct, lacking the C-terminus, showed subcellular localization similar to hDVL1^1519ΔT^ (E) Notably, both hDVL1^1519ΔT^ and hDVL1^1431*^ led to a significantly increased nuclear accumulation compared to wt hDVL1, as determined by Two-way ANOVA with Tukey’s post hoc test. (F) The fraction of total (%) for subcellular localization of all hDVL1 constructs. Key: *p < 0.05, **p < 0.01, ***p < 0.001, ****p < 0.0001, ns – not significant

**Figure 8.**
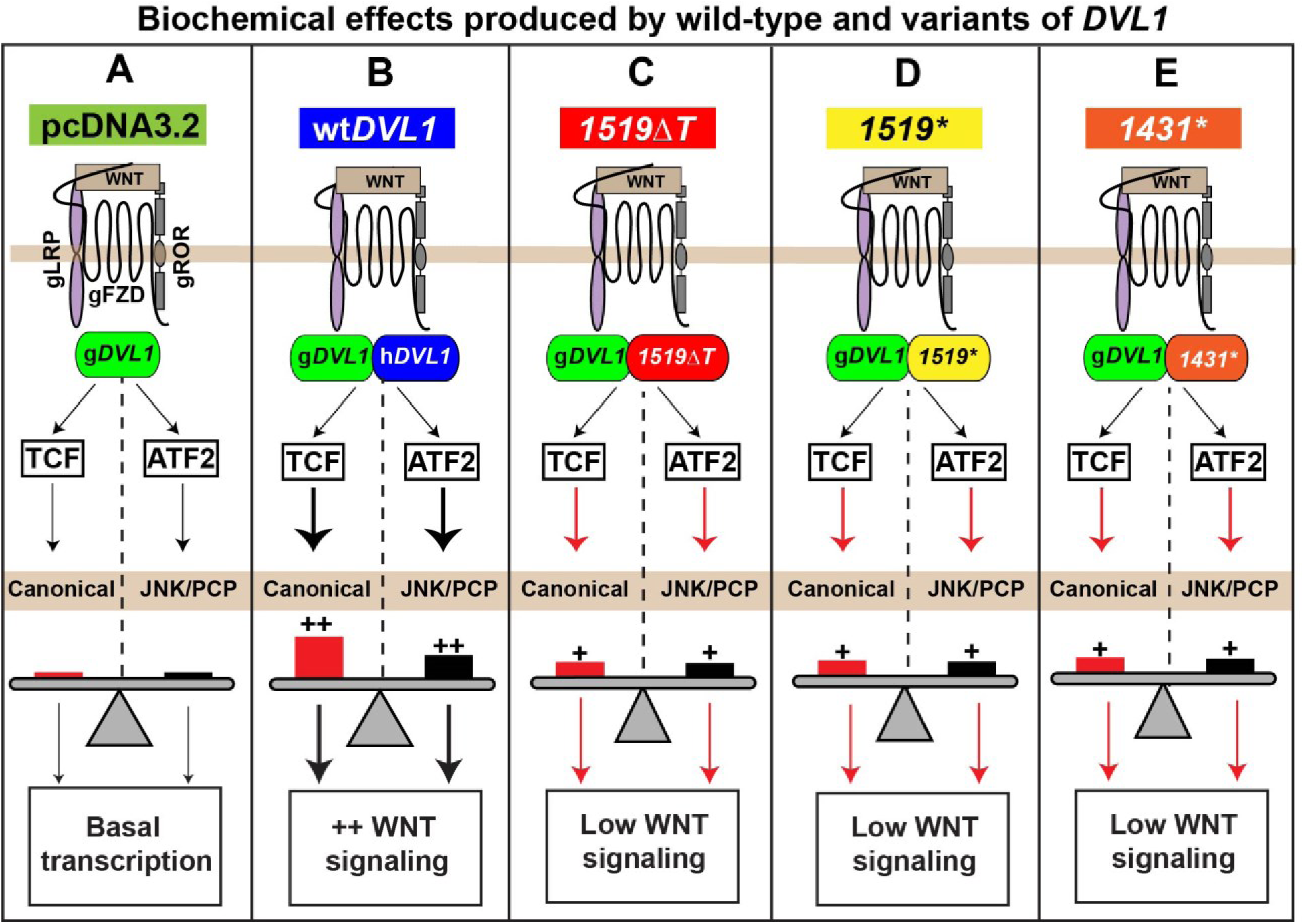
Summary of biochemical signaling effects produced by wt h*DVL1* and three h*DVL1* variants in frontonasal mass mesenchymal cells. The activity of plasmids measured as fold-change compared to empty plasmid is summarized schematically. (A) Overexpression of the control/parent plasmid (pcDNA3.2) in frontonasal mass mesenchyme (have endogenous DVL1; green), showed basal level activity in both the canonical (STF, red) and non-canonical JNK-PCP WNT pathways (ATF2, black). (B) Overexpression of wt h*DVL1* (blue) significantly activated canonical and non-canonical reporters. (C) The h*DVL1^1519^*^Δ*T*^ (red), (D) h*DVL1^1519*^* (yellow), (E) and h*DVL1^1431*^* (orange) variants weakly activated both STF and ATF2 compared to wild-type h*DVL1* but the activity was above the basal level.

## Discussion

In this study we investigated human mutations in *DVL1* that cause Robinow syndrome. Every *DVL1* mutation associated with ADRS clusters within the penultimate exon and the last exon, causing a frameshift in the sequence and subsequent translation into an abnormal peptide at the C-terminus (Hu et al., 2022; White et al., 2015; Zhang et al., 2022). The novel C-terminal peptide has no known homology and is conserved among individuals with the *DVL1* form of Robinow syndrome (White et al., 2015; Zhang et al., 2022). In individuals with *DVL3* frameshift mutations a different C-terminal peptide is generated (White et al., 2016). There are no reported human DVL1 variants that affect the main functional domains (DIX, PDZ, DEP) which suggests that such mutations are incompatible with life. Our study of ADRS frameshift mutations in *DVL1* has shed light on the pathogenesis of the syndrome and on the requirements of the *DVL1* C-terminus in development, skeletal differentiation and cell signaling.

### Connections between the ARDS-*hDVL1* frameshifts, chicken facial morphogenesis and the facial phenotypes in RS

All RS individuals have a wider nasal bridge, hypertelorism, frontal bossing and maxillary hypoplasia (Beiraghi et al., 2011; Conlon et al., 2021; Mazzeu and Brunner, 2020). Some of the phenotypes produced in chicken embryos can explain the origins of theses specific RS dysmorphologies. We found that the frontonasal mass width was increased by h*DVL1* variant. These effects are unrelated to neural crest cell migration since viruses are injected after neural crest cells have settled into the face. We also ruled out increased proliferation as the mechanism contributing to facial widening. We also showed that overexpression of human *DVL1* whether wt or mutant had no effect on endogenous genes. This suggested that the mutant variants may have prevented the normal process of narrowing of the frontonasal mass (Danescu et al., 2021). The midfacial narrowing via convergent-extension morphogenesis also occurs in human embryos (Diewert and Lozanoff, 1993). The mechanism of narrowing is related to cytoskeletal arrangement regulated by the RhoA-GTPase pathway. A previous study from our lab where ROCK (the direct target of RhoA) was inhibited, the frontonasal mass failed to narrow (Danescu et al., 2021). Since h*DVL1* signals via DAAM to activate the RhoA and Rock pathway (Habas et al., 2001), it is possible that this branch of WNT signaling is less active in the presence of the variant form of h*DVL1*. Indeed, ATF2 luciferase was less active in the presence of h*DVL1^1519^*^Δ*T*^ compared to wt h*DVL1*.

Other facial phenotypes produced by the h*DVL1* overexpression in chicken embryos such as the shorter beak and curvature towards the injected side recapitulates midface hypoplasia observed in patients with ADRS (Bunn et al., 2015; Conlon et al., 2021). Indeed, the h*DVL1^1519^*^Δ*T*^ variant caused significant shortening and deviation of the upper beak with a 92% penetrance. It is possible that the decreased proliferation we identified in stage 29 embryos could explain the decreased length of the upper beak. Other mechanisms underlying jaw deficiencies seen in RS could be a failure of membranous bone differentiation. In the chicken embryo as with all vertebrates, the facial bones such as premaxilla, maxilla and mandible are formed by direct ossification of the neural crest-derived mesenchyme (Ornitz and Marie, 2002). There is no cartilage precursor for intramembranous bone, therefore the jaw hypoplasia in RS patients is due either to smaller condensations in the early stages or decreased apposition of bone in later stages of bone remodeling. In our study the premaxilla failed to form in 12/13 embryos injected with *hDVL1^1519^*^Δ*T*^ virus. This suggests that the *hDVL1^1519^*^Δ*T*^ variant interferes with the early mesenchymal condensation stages. Verification of this hypothesis requires a marker specific for intramembranous bone condensations.

### The effects of *hDVL1* on canonical WNT signaling vary according to context

We showed that the h*DVL1^1519^*^Δ*T*^ variant was significantly less able to activate the canonical luciferase reporter (STF) in primary facial mesenchyme compared to wt *hDVL1*. The same h*DVL1* variant was transfected into HEK293 cells by our group and a similar result was observed (Gignac et al., 2023). In that study we also showed that the equimolar combination of variant h*DVL1* and wt h*DVL1* led to a dominant negative inhibition of the wt protein in luciferase assays. Thus, in our study the lower activation could also be partly due to dominant negative effects of variant h*DVL1* on activity of g*DVL1*. The exact reason why variant h*DVL1* inhibits canonical WNT signaling is unclear. It is possible that the abnormal peptide removes an important binding site for a protein that is needed for canonical WNT signal transduction. For example, IQGAP1 (Ras GTPase-activating-like protein) associates with the C-terminus of DVL1 and has been shown be essential to activate β-catenin mediated canonical WNT activity (Goto et al., 2013). We hypothesize that the loss of activity in the canonical pathway may be attributed loss of interaction of mutant h*DVL1* with a protein such as IQGAP1.

We looked back at other assays carried out with h*DVL1^1519^*^Δ*T*^ to gain an overview of how this variant affects canonical WNT signaling. Nuclear migration of β-catenin and displacement of Groucho is required for increased transcriptional activity (Daniels and Weis, 2005; Vuong and Mlodzik, 2022). We expected to see reduced nuclear β-catenin in vivo in the frontonasal mass presence of variant *hDVL1* but there were no changes in the proportion of positive cells. The levels of total g*CTNNB1* were measured, 48h after virus infection in vivo. However, the RNA levels of g*CTNNB1* were also not changed. Additionally, we also did not see any effects on nuclear β-catenin levels in western blot analysis. The qRT-PCR experiments of micromass culture collected at day 8 also showed no significant changes in g*CTNNB1* with the ADRS variant. To test if the h*DVL1^1519^*^Δ*T*^ affects nuclear transport of β-catenin, we should examine the expression of nuclear β-catenin protein or g*CTNNB1* RNA in micromass cultures at earlier stages to match the luciferase assays. Taken together our data suggests the h*DVL1^1519^*^Δ*T*^ transiently inhibits canonical WNT signaling during critical stages when intramembranous bone condensations are developing.

We examined several bone and cartilage markers in vivo, 48h after virus infection with the h*DVL1^1519^*^Δ*T*^ virus. There were no significant differences in *RUNX2* or *MSX1* transcript levels, markers that typically are expressed during intramembranous bone formation. Similarly, in micromass cultures, there was no significant difference in *MSX1* expression. Therefore, to verify the hypothesis that intramembranous bone progenitor cells are reduced by the ADRS h*DVL1* variant, another expression experiment would need to be carried out on two-day micromass cultures to better match the timing of the luciferase experiments. In addition, a full RNA-seq experiment on the chicken embryos injected with virus would provide broad insights into the full transcriptome response to the h*DVL1* constructs.

### What did we learn about C-terminus function in RS and normal development?

One of the most unexpected findings was that loss of almost all of the hDVL1 C-terminus h*DVL1^1519*^*(DVL1^507*^) permits normal development. This suggests that the looping back of the most terminal part of the C-terminus to bind to a PDZ binding site (autoinhibition) (Itoh et al., 2005; Qi et al., 2017), may not critical for the function of *DVL1* during development in chicken embryos. The novel finding here is that first 30 amino acids of the C-terminus of hDVL1, retained in DVL1^507*^, are important to maintain normal function. Indeed, nuclear and cytoplasmic distribution of DVL1^507*^ were identical to wt hDVL1. This suggests that the NES of hDVL1 is situated within these 30 amino acids. The only evidence that this DVL1^507*^ form of hDVL1 is not behaving the same as wt h*DVL1* is in two experiments - the in vivo qRT-PCR experiments and in the luciferase assays. In the qRT-PCR, the h*DVL1^1519*^* significantly decreases g*CTNNB1* expression. In luciferase assays, the h*DVL1^1519*^* behaves similarly to the ADRS variant and weakly activates STF and ATF2. This suggests that even though the truncated form, DVL1^507*^, may be sufficient for craniofacial development, it has lost the components critical for WNT signaling.

The phenotypes caused by overexpression of the h*DVL1^1431*^*construct were severe compared to h*DVL1^1519^*^Δ*T*^, since other organs were affected and there was high lethality. As far as we know, there are no reported human truncating mutations that affect the C-terminus of hDVL1 (ncbi.nlm.nih.gov/ClinVar). All h*DVL1* constructs in this study retain the nuclear NLS located between the PDZ and DEP. Consequently, all forms of hDVL1, whether wt, mutant or truncated are able to enter into the nucleus. Based on the predicted location (Itoh et al., 2005; Pruller et al., 2022; Sharma et al., 2018), NES absent in the DVL1^1431*^. Therefore, 80% DVL1^1431*^ protein is exclusively localized in the nucleus. Others have shown that hDVL1 missing the N-terminal part of the DEP domain was strictly localized in the nucleus (Paclikova et al., 2017). Surprisingly, 80% of the hDVL1^1519ΔT^ variant protein was also nuclear. This is the first data to suggest that the abnormal C-terminus may be influencing the phenotype, potentially owing to an abnormal tertiary structure. Given that the aberrant C-terminus is found in all 18 frame-shifted h*DVL1* variants, it is thus playing a role in patient phenotypes.

### Materials and methods Chicken embryo model

White leghorn eggs (*Gallus Gallus*) obtained from the University of Alberta were incubated to the appropriate embryonic stages, based on the Hamilton Hamburger staging guide (Hamburger and Hamilton, 1992). All experiments were performed on prehatching chicken embryos which are exempted from ethical approval by the University of British Columbia Animal Care Committee and the Canadian Council on Animal Care.

### Cloning of human wt and variant human *DVL1* constructs

The open reading frame encoding human DVL1 was purchased from Origene (#RC217691). The insert was initially in a pCMV6 vector containing a full coding sequence with two C-terminal tags: myc and DDK (Flag similar). To change pCMV6 vector to an entry clone (pDONR221), gateway cloning with BP clonase II (ThermoFisher #11789020) was performed. Then restriction free cloning was used to insert kozak-N-terminal flag (DYKDDDDK) sequence and 3’ C-terminal STOP (added to eliminate commercial tags from company) to this pDONR221 vector. Site-directed mutagenesis was performed in the pDONR221 (Gateway compatible) vector and then recombined into destination vectors (pcDNA3.2 expression vector and RCAS retroviral vector) using LR Clonase II enzyme (ThermoFisher # 11791019). Gateway cloning (ThermoFisher) was used to move human *DVL1* into the pDONR221. Site-directed mutagenesis was used to knock-in the mutations. Autosomal dominant RS *DVL1* frameshift mutation (OMIM: 616331) was knocked in (*1519*Δ*T*) (White et al., 2015). Two additional h*DVL1* constructs were made for this study - either a STOP codon was placed immediately after nucleotide 1519 (*DVL1^1519*^*, DVL1^507*^) or the entire C-terminus was prevented from being expressed by placing a stop codon after the DEP domain (*DVL1^1431*^*, DVL1^477*^). To create the RCAS viruses, a Gateway compatible RCASBPY destination vector was used for recombination with the pDONR221 vector using LR Clonase II (Loftus et al., 2001). The location of *DVL1* containing viruses was determined retrospectively on histological sections stained with a Group-associated antigens (GAG), an antibody that recognizes specific proteins in the RCAS virus.

### Growth of RCAS viral particles and viral titre

Replication-Competent ASLV long terminal repeat (LTR) with a Splice acceptor (RCAS) (Hughes, 2004; Loftus et al., 2001) plasmid DNAs (2.5µg) encoding *GFP*, human *DVL1*, or three *DVL1* variants were transfected into the DF-1 immortalized chicken fibroblast cell line (American Type Culture Collection, CRL-12203) using Lipofectamine 3000 (Thermo Fisher #L3000-008) following the manufacturer’s guidelines. RCAS virus containing *GFP* insert (kindly provided by A. Gaunt) served as a control in virus overexpression studies, as published(Geetha-Loganathan et al., 2014; Gignac et al., 2019; Gignac et al., 2023; Hosseini-Farahabadi et al., 2013; Hosseini-Farahabadi et al., 2017). DF1 cells were cultured at 37°C and 5% CO_2_ in DMEM (ThermoFisher#1967497) medium supplemented with 10% fetal bovine serum (FBS; Sigma #F1051) and 1% penicillin/streptomycin (ThermoFisher #15070-063). The cells were maintained in 100 mm culture dishes with media changes every other day and passaged 1:2 two to three times per week using trypsin-EDTA (0.25%, ThermoFisher #25200-072). After six weeks of culturing, the viral particles were collected and centrifuged in a swing bucket SW28 rotor (Beckman #97U 9661 ultracentrifuge) for 2.5 hours (no brake) at 25,000 rpm at 4°C. The supernatant was carefully removed, and the resulting pellet was resuspended in 50-100 µl Opti-MEM (ThermoFisher#319850962). This suspension was then incubated overnight at 4°C. The concentrated viral particles obtained were aliquoted (5µL aliquots), rapidly frozen in methanol + dry ice, and stored at −80°C for future use (Goodnough et al., 2012).

To determine viral titer, 50-60% confluent DF1 fibroblasts were infected with serial dilutions of 2μl of concentrated viral stock. After 36 hours of virus incubation, cells were fixed in 4% paraformaldehyde for 30 mins. Immunocytochemistry with Group Associated Antigens (GAG) antibody was performed on virus-treated cells (Table S9). The cells were permeabilized for 30 mins with 0.1% Triton X-100, followed by blocking in 10% goat serum and 0.1% Triton X-100 and overnight incubation with the primary antibody (Table S9). Fluorescence images were captured using a Leica inverted microscope at 10x with a DFC7000 camera. The analysis of viral titer was done with ImageJ’s cell counter tool by determining the proportion of GAG-positive cells per mL in a 35mm culture plate. Virus titer = # of GAG positive cells * (Area counted * total cells expressing GAG)/2*1000 (Fig. S7).

### Chicken embryo injections

Fertilized eggs obtained from the University of Alberta, Edmonton, were incubated in a humified incubator at 38°C until Hamilton Hamburger (Hamburger and Hamilton, 1951; Hamburger and Hamilton, 1992) stage 15 (E2.5). Concentrated RCAS (titer = >2 x 10^8^ IU/mL) retrovirus viral particles (∼5μl) combined with Fast Green FCF stain (0.42%, Sigma #F7252) (1μl) were injected into the frontonasal mass (anatomic region bounded by the nasal slits) of stage 14-15 chicken embryos (25-28 somites) using glass filament needles (thin-wall borosilicate capillary glass with microfilament, A-M systems #615000) and a Picospritzer microinjector (General valve corp. #42311). The infection of embryos with RCAS at stage 15 (E2.5) was performed to ensure maximum infection of facial prominences(Geetha-Loganathan et al., 2009; Hosseini-Farahabadi et al., 2017). Due to accessibility, all injections were made into the right frontonasal mass as the chick embryos turn on their left side during development. The facial prominences form around stage 20, and the complex and temporally regulated patterning occurs between stages 20-29. The skeletal derivatives of the frontonasal mass are fully patterned and ossified between stages 34-40. The investigation encompassed multiple embryonic stages to comprehensively analyze these developmental processes. After conducting the overexpression of high-titer *DVL1* viruses at stage 15, the retrospective determination of virus location was carried out on histological sections. These sections were stained with Group-associated antigens (GAG), an antibody that identifies specific proteins in the RCAS virus (Geetha-Loganathan et al., 2009; Gignac et al., 2023; Hosseini-Farahabadi et al., 2017) (Table S9).

### Wholemount staining of skulls

To study skeletal elements, embryos were grown until stage 38 (10 days post-injection, Table S3). The embryos were washed in 1x phosphate-buffered saline (PBS; 137 mM NaCl, 8.1 mM Na_2_HPO_4_, 2.7 mM KCl, 1.5 mM KH_2_PO_4_; pH 7.3) and fixed in 100% ethanol for 4 days. After removal of eyes and skin, the embryos were transferred to 100% acetone for another 4 days. Subsequently, the heads were stained with a freshly prepared bone and cartilage stain (0.3% Alcian blue 8GX (Sigma #A5268) in 70% ethanol, and 0.1% alizarin red S (Sigma # A5533) in 95% ethanol, with 1 volume of 100% acetic acid and 17 volumes of 70% ethanol) for two weeks on shaker at room temperature. Following staining, the skulls were washed in water and cleared in a 2% KOH/20% glycerol solution on shaker for 4 days, followed by immersion in 50% glycerol for imaging. The heads were stored in 100% glycerol post-imaging. Phenotyping was conducted by photographing the right lateral, superior and palatal views of cleared heads using a Leica DFC7000T microscope camera. Skeletal preparations from each virus type were analyzed for changes in the size or shape of bones derived from the frontonasal mass, missing bones, or qualitative reduction in ossification observed as reduced alizarin red stain. Statistical analysis was performed using contingency analysis Chi-square test in GraphPad Prism 10.1.0.

### Primary cultures of frontonasal mass mesenchyme

Stage 24 chicken embryos were extracted from the eggs and their extra-embryonic tissues were removed in cold phosphate-buffered saline (PBS). The frontonasal mass was dissected in cold Hank’s balanced saline solution (HBSS) (without calcium and magnesium) (ThermoFisher #14185052) with 10% FBS and 1% Antibiotic-Antimycotic (Life Technologies #15240-062). Dissected frontonasal pieces were incubated in 2% trypsin (Gibco) at 4°C for 1 hour. Hank’s solution was added to inhibit the enzymatic activity of trypsin. Ectoderm was manually peeled off from the frontonasal mass pieces. The cell solution was then centrifuged at 1000 g, 4°C for 5 minutes. The supernatant was removed and the frontonasal mass pieces were resuspended in Hank’s solution. The mesenchymal cells were counted using a hemocytometer and 2×107 cells/ml were resuspended in chondrogenic media (micromass media) containing DMEM/F12 medium (Corning #10-092-CV) supplemented with 10% FBS, 1% L-Glutamine (ThermoFisher #25030), Ascorbic acid (50 mg/ml) (ThermoFisher #850-3080IM), 10 mM β-glycerol phosphate (Sigma Aldrich #G9422) and 1%, Antibiotic-antimycotic (ThermoFisher #15240-062). The cells in suspension were subsequently infected with 3µL *GFP* (control), wt h*DVL1*, or h*DVL1* variants containing viruses. The 10µl of cells suspension infected with virus was plate micromass cultures (3-4 spots per 35mm culture dish, NUNC #150318) at a density of 2 x 10^7^ cells/ml, (Hosseini-Farahabadi et al., 2013; Richman and Tickle, 1989; Richman and Tickle, 1992; Underhill et al., 2014). The culture plates were incubated at 37 °C and 5% CO_2_ for 90 minutes to allow cells to attach and then flooded with 2 ml of micromass media. Thereafter, micromass culture media was changed every other day for experimental time points of day 4, 6, and 8.

### Wholemount staining of micromass cultures

On day 4, 6, and 8, cultures were fixed in 4% paraformaldehyde for 30 minutes at room temperature and subjected to wholemount staining. To detect the mineralization of cartilage using alkaline phosphatase stain (Table S8), fixed cultures were incubated at room temperature in 100mM Tris for 30 minutes (pH 8.3). Following this, the cultures were stained with 0.5% Alcian Blue in 95% EtOH: 0.1 M HCl (1:4) to detect the area occupied by cartilage, as previously described (Hosseini-Farahabadi, 2013; Underhill, 2014). All cultures were counterstained with 50% Shandon’s Instant Hematoxylin (ThermoFisher #6765015). The stained cultures were photographed under standard illumination using a stereomicroscope (Leica #M125). Wholemount staining was conducted on three biological and three technical replicates, and the experiment was repeated five times.

### Histology and Immunofluorescence

Embryos collected at stage 28 (Table S5) or micromass cultures (day 4, 6, 8, Table S7) were fixed in 4% PFA. The embryo samples were immersed in the fixative for 2-3 days at 4°C. The RCAS-infected cultures fixed in 4% paraformaldehyde for 30 mins. The cultures were then removed from the plate using a cell scraper (ThermoFisher #08-100-241), embedded in 2% agarose (Sigma #A9539) on a cold ice slab, and subsequently wax-embedded. The embryos (positioned frontally) and cultures (positioned transversely) were embedded in paraffin wax and sliced into 7µm sections. The sections were then utilized for histological and immunostaining analysis.

Selected frontal (embryos) and transverse (micromass cultures) sections were stained to visualize the differentiated cartilage and bone. Sections were dewaxed in xylene, rehydrated from 100% ethanol to water, and stained with 1% Alcian blue 8GX (in 1% acetic acid) for 30 minutes. After staining, sections were rinsed in 1% acetic acid and water. Subsequently, sections were stained in Picrosirius Red (0.1% Sirius Red F3B in saturated picric acid) for 1 hour in dark, followed by rinsing in 1% acetic acid, dehydration through ethanol, back to xylene, and mounted with Shandon Consul-mount (Thermo Scientific #9990441).

Immunofluorescence analysis was conducted on in vivo and day 6 and 8 cultures. Specific antibodies and treatments performed for each assay are outlined in Table S9. Primary antibodies were allowed to incubate overnight at 4°C, while secondary antibodies were incubated at room temperature for 1.5 hours unless otherwise specified. Sections were counterstained with Hoechst (10μg/ml #33568, Sigma) and incubated for 30 minutes at room temperature, then mounted with Prolong Gold antifade (ThermoFisher #P36930). Fluorescence images were captured using 20X objective on a slide scanner (3DHISTECH Ltd., Budapest, Hungary).

### Apoptosis and Cell proliferation

Apoptosis was analyzed using TUNEL (Terminal deoxynucleotidyl transferase dUTP nick end labeling) assay on sections obtained from virus infected frontonasal mass stage28 and micromass cultures sections day 6 and day 8. The TUNEL assay was performed using ApopTag Plus *in Situ* Apoptosis Fluorescein Detection Kit (Millipore Sigma # S7111).

For cell proliferation studies, embryos at stage 28, stage 29 or stage 30 were labeled with 50µl of 10 mM BrdU (Bromodeoxyuridine; Sigma #B5002) and incubated at 38°C for 1 hour prior to euthanizing. For labeling micromass cultures, 50µl of 10mM BrdU was added to the culture media (37°C 5% CO_2_) 1 hour before fixing day 6 and day 8 cultures in 4% PFA. Immunostaining was performed on the sections with anti-BrdU (Developmental Studies Hybridoma bank, 1:20, #G3G4) as described in Table S9. Fluorescence images were collected with a 20X objective on a slide scanner (3DHISTECH Ltd., Budapest, Hungary).

### Immunocytochemistry with transfected human *DVL1* variants

HEK293T DVL Triple Knockout (TKO) cells were received as a kind gift from Dr. Stephane Angers (University of Toronto) (Cervenka et al., 2016). Cells were grown to 60% confluency and transfected using Lipofectamine3000 (ThermoFisher #L3000-008) following manufacturer’s instructions. Transfected plasmids include (2.5 μg DNA): parent plasmid pcDNA3.2, wt*DVL1, DVL1^1519ΔT^, DVL1^1519*^, DVL1^1431*^*. Post transfection, the cells were grown for 48h and then fixed in 4% PFA for 20 min and then stored in 1X PBS at 4°C overnight. Cultures were blocked in 10% normal goat serum with 0.2% triton-X in 1 X PBS for 1h then incubated overnight at 4°C with primary anti-Flag antibody. Secondary antibody was applied for 1h at room temperature and counterstained with 10 µg/ml Hoechst (Table S8). Cells were imaged using 20x dry objective Leica THUNDER 3D cell imager widefield microscope, Leica K8 camera.

### qRT-PCR on frontonasal mass in vivo and in vitro

Viral spread in the frontonasal mass was quantified using primers specific to human *DVL1* (primer set, Table S10). Three biological replicates containing five to six pieces of the right half of the frontonasal mass pooled in each sample were harvested for each virus at 48 h post injection. Similarly, three biological replicates containing pools of twelve micromass cultures per replicate were collected on day 6 and day 8. Total RNA was isolated from frontonasal masses using Qiagen RNAeasy kit (#75144, Toronto, Canada). Sybr green-based quantitive reverse transcriptase polymerase change reaction (Advanced universal SYBR® Green supermix; Bio-Rad #1725271) (qRT-PCR) was carried out using an Applied Biosystems StepOnePlus instrument. qRT-PCR cycling conditions were - 95°C for 10min, 40X (95°C for 5s, 60°C for 20 seconds). Analysis used human-specific primers for *DVL1* and avian primers were used (Table S9). The expression of each biological replicate was normalized to 18s RNA (Applied Biosystems, 4328839) and then these ΔCt values were used to calculate the ΔΔCt relative to the average levels of expression of the gene in *GFP*-infected cultures. The ΔΔCt method was used to calculate relative fold-change expression between RS-*DVL1* infected frontonasal mass and *GFP* as described (Schmittgen and Livak, 2008). Statistical analysis was done with one-way ANOVA Tukey’s post hoc test in GraphPad Prism 10.0.2. A sample size calculator was used to determine how many samples would need to be included in order to detect a P value of 0.05 80% of the time and it was necessary to collect 13 biological replicates. This number of biological replicates was not feasible for these studies.

### Luciferase reporter assays

Transient transfections for luciferase assays were performed in untreated stage 24 frontonasal mass mesenchymal micromass cultures (Geetha-Loganathan et al., 2014; Hosseini-Farahabadi et al., 2013). Cells were transfected with Lipofectamine 3000 (Invitrogen, L3000-008, Nunc 24-well plates #142475) transfection reagent. Micromass cultures were allowed to attach for 45 mins after plating and transfection reagents were added to the culture spot 30 mins prior to flooding the culture plate with micromass media. The following plasmids were used: control/empty (pcDNA3.2/V5-DEST), h*DVL1*, *DVL1^1519^*^Δ*T*^, *DVL1^1519*^*, *DVL1^1431*^*. Firefly reporter plasmids: SuperTopflash (STF; 0.2ug, Addgene plasmid #12456) and Activating Transcription Factor 2 (0.4ug; ATF2) (Ohkawara and Niehrs, 2011) along with Renilla luciferase was transfected for normalization (0.01μg). Renilla luciferase was used as normalization control. Assay reading was done 48h after transfection representing day 3 of culture. The dual-luciferase reporter assay system (Promega #E1910) was used for all luciferase assays as described (Geetha-Loganathan et al., 2014). Luminescence activity was detected with a PerkinElmer Victor X2 Multilabel Microplate Reader at 1 s reading with OD1 filter. All data shown represents two to three independent experiments with three technical and three biological replicates carried out for each transfection mixture. Statistical analysis done using one-way ANOVA, Tukey’s post hoc test in GraphPad Prism 10.0.2. The number of biological replicates was determined by our previous studies using luciferase assays (Gignac et al., 2019; Gignac et al., 2023).

### Image analysis and statistics

1. For measurement of the width of the frontonasal mass at stage 29, distance between the nasal slits (illustrated in Fig. 3A) was measured manually with linear measurement annotation tool in CaseViewer (version 2.4). The data was analyzed using one-way ANOVA, Tukey’s test in GraphPad Prism 10.1.0.
2. The thickness of 8 micromass cultures was measured using histological sections stained with Alcian blue and picrosirius red. The linear measurement annotation tool in CaseViewer (version 2.4) was utilized. For each genotype, measurements were performed on three biological replicates. Each biological replication represents an average of three ribbons. The data was analyzed using one-way ANOVA, Tukey’s test in GraphPad Prism 10.1.0.
3. For analysis of immunofluorescence staining performed on stage 29 samples, cells in the prenasal cartilage in all treated samples were counted in a 100 x 100µm^2^ area to get the average cell density. The right frontonasal mass was divided into four regions (100 x 250um^2^) (Fig. 2U) to count the proportion of cells expressing BrdU (n=3) and β-catenin (n=3). All cell counts were performed twice with the counter plugin in ImageJ by a blind observer. The data was analyzed using one-way ANOVA, Tukey’s test in GraphPad Prism 10.1.0. Similar samples sizes were used in other studies on BrdU labelling (Gignac et al., 2019; Gignac et al., 2023).
4. For analysis of subcellular location of Flag-DVL1 protein in HEK293T DVL TKO cells, 150 transfected cells in random fields of view were counted in three biological replicates (three culture plates) (450 cells per virus type). The expression of FLAG was classified as cytoplasmic, nuclear, cytoplasmic + nuclear, and DVL1 forming puncta. Statistical analysis was conducted using two-way ANOVA, Tukey’s post hoc test and contingency analysis Chi-square test in GraphPad Prism 10.1.0.

## Acknowledgements

We are grateful to the Developmental Studies Hybridoma Bank (IA, USA) for providing antibodies. We thank Dr. Janel Kopp for use of the 3DHistech slide scanner for all chicken histology images. We acknowledge the excellent virus injections carried out by Julian Kim and Adrian Danescu.

## Author contributions

Conceptualization: S.S.T and J.M.R.; Methodology: S.S.T and K.F.; Formal analysis: S.S.T. and J.M.R. Investigation: S.S.T., K.F., J.M.R.; Resources: J.M.R., E.M.V.; Writing - original draft: S.S.T. and J.M.R.; Writing - review & editing: S.S.T., J.M.R, E.M.V; Supervision: J.M.R.; Project administration: J.M.R., E.M.V.; Funding acquisition: E.M.V., J.M.R.

## Data availability

All relevant data can be found within the article and its supplementary information. The senior author will provide additional data and DNA constructs on request.

