## Supplementary material for "*DVL1* variants and C-terminal deletions have differential effects on craniofacial development and WNT signaling": Tophkhane et al. Supplementary Data (from another version of this preprint: version 1, 2024-02-29)

### Supplementary figures

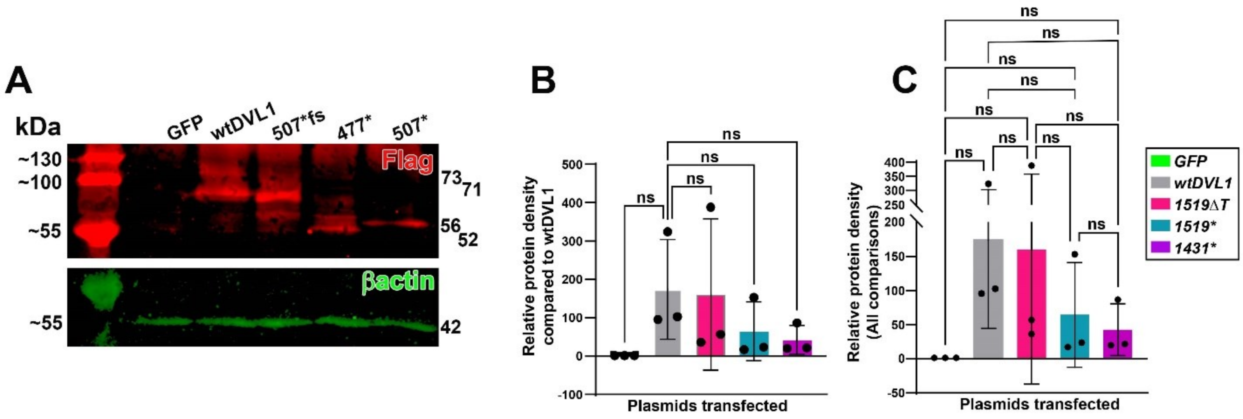


**Figure S1: Western blot analysis was conducted on DF1 chicken fibroblast lysate.**

The cells were transfected with FLAG-tagged hDVL1 construct and cultured for 4 weeks before lysis. (A) The Western blot probed with anti-FLAG antibody (Sigma, #F7425) shows comparable hDVL1 protein levels. Notably, there are expected size variations among the DVL1 proteins shown on the right side of the blot, wild-type DVL1 (~73 kDa), DVL1$^{507fs*142}$ (~71 kDa), DVL1$^{507*}$(~56 kDa), and DVL1$^{477*}$(~52 kDa). β-actin (Abcam 1:2000 #8226, 42kDa) was used as a loading control. (B) Graph showing quantification of relative protein density compared to wtDVL1 (One-way ANOVA, Dunnett's test), and (C) all comparisons (One-way ANOVA, Tukey's text), (n=3 independent blots). Statistical analysis was done in GraphPad Prism 10.1.0. Key: n= non-significant.

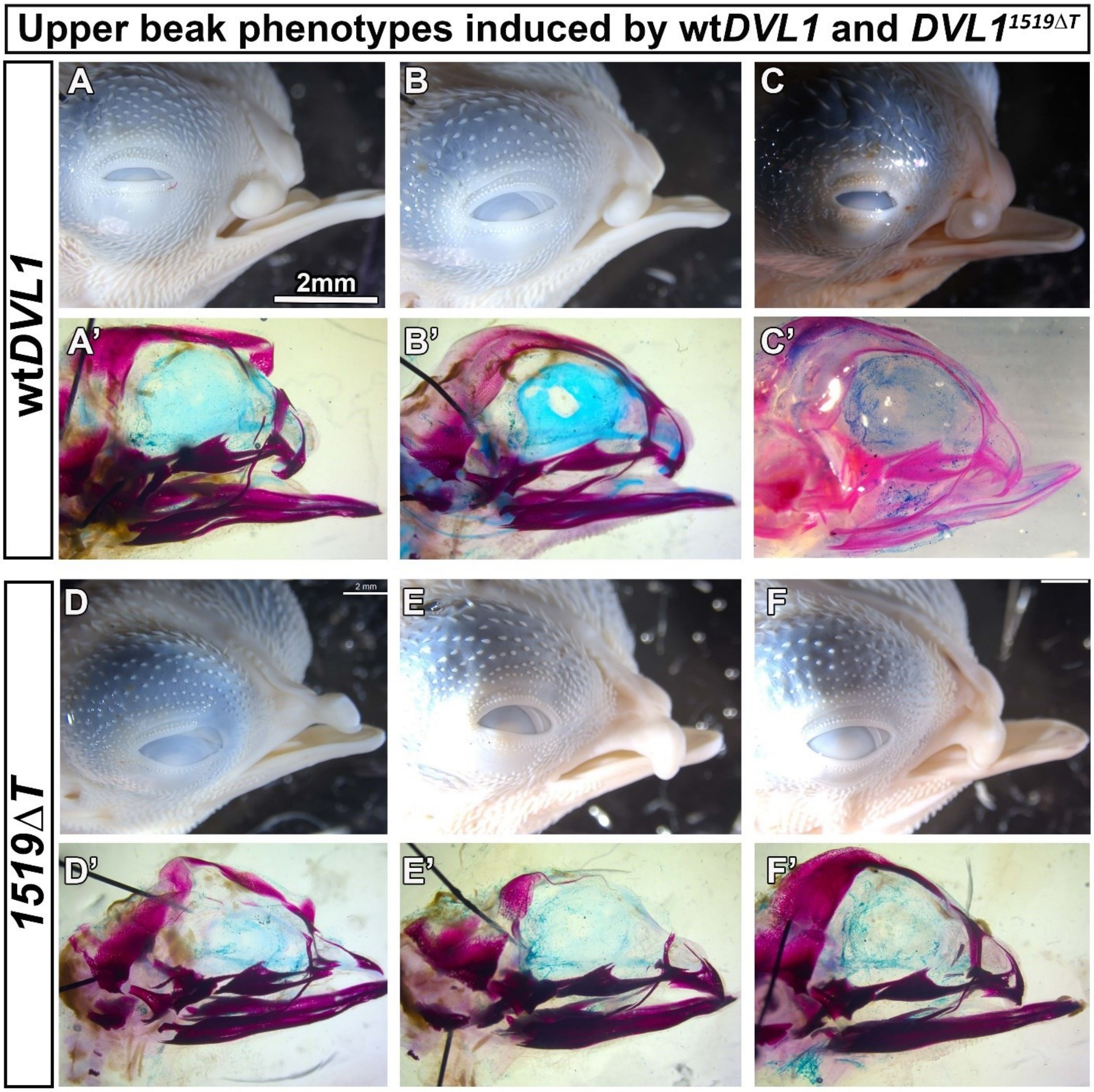


**Figure S2: Additional specimens treated with RCAS: wild-type hDVL1 and RCAS:hDVL1$^{1519\Delta T}$**

Embryos injected with RCAS wild-type *DVL1* or *DVL1$^{1519\Delta T}$* variant into the right frontonasal prominence at stage 15 (E2.5) and fixed 10 days post injection at stage 38 (E12.5). (A-C, D-F) External photographs in right lateral view of stage 38 embryos showing shortening and deviation of the upper beak caused by wild-type or mutant h*DVL1* viruses. (A'-C', D'-F') Wholemount skulls stained with alcian blue (for cartilage) and alizarin red (for bone) for the same specimens showing abnormal development of the skeletal elements. The faint cartilage stain (blue) in all skulls is a technical error. Scale bar: A-F' = 2mm.

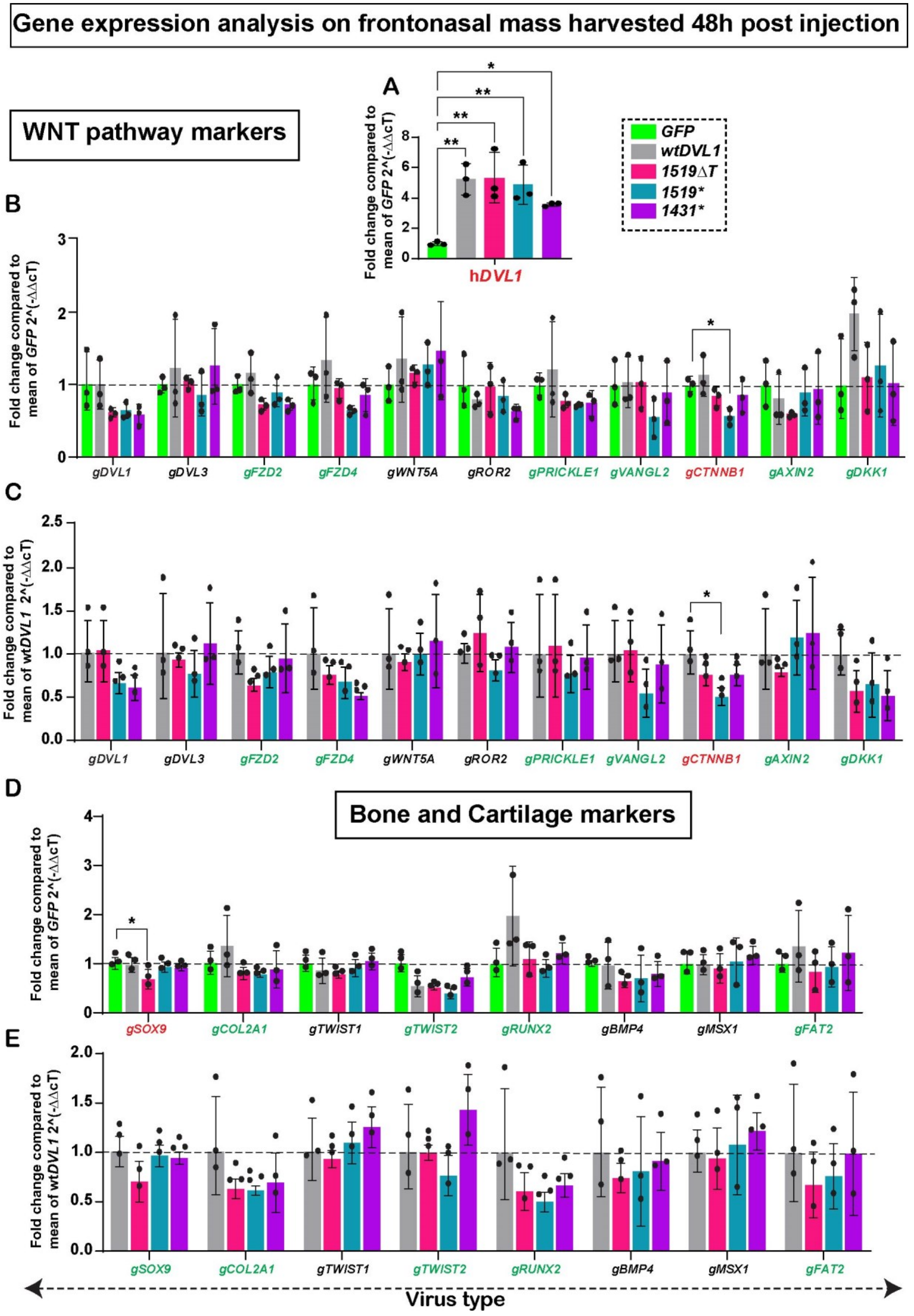
Gene expression analysis on frontonasal mass harvested 48h post injection
A
WNT pathway markers
Bone and Cartilage markers
GFP
wtDVL1
1519ΔT
1519*
1431*
hDVL1
Fold change compared to mean of GFP 2^(-ΔΔcT)
Fold change compared to mean of wtDVL1 2^(-ΔΔcT)
B
C
D
E
gDVL1
gDVL3
gFZD2
gFZD4
gWNT5A
gROR2
gPRICKLE1
gVANGL2
gCTNNB1
gAXIN2
gDKK1
gSOX9
gCOL2A1
gTWIST1
gTWIST2
gRUNX2
gBMP4
gMSX1
gFAT2
Virus type

**Figure S3: Gene expression analysis in frontonasal mass infected with hDVL1 viruses harvested 48h post treatment**

qRT-PCR analysis of RNA isolated from the virus infected right half of the frontonasal mass harvested at 48h post-infection with either *GFP* (control), wild-type h*DVL1* or h*DVL1* variants. The expression of each biological replicate was normalized to 18s RNA and then these ΔCt values were used to calculate the ΔΔCt relative to the average levels of expression of the gene in *GFP*- or wild-type h*DVL1* infected cultures. The level of expression of h*DVL1* was measured with human-specific primers. (A) There was a 4-5-fold higher expression of h*DVL1* in all h*DVL1*-infected culture conditions in comparison to *GFP* virus. *Gallus* specific primers were used for WNT, BMP and chondrogenic markers. Statistical comparisons were made with (B, D) *GFP* or (C, E) wild-type h*DVL1*. (B, C) There was no change in the expression of g*DVL1* and g*DVL3* induced by the h*DVL1* viruses). (B, C) Significant decreases in expression of g*CTNNB1* was caused only with h$DVL1^{1519*}$ virus compared to both *GFP* and wild-type h*DVL1*. (D) The expression of g*SOX9* was downregulated by $DVL1^{1519\Delta T}$ compared to *GFP*. (B-E) The expression other candidate genes (green text) including g*FZD2,* g*FZD4,* g*PRICKLE1,* g*VANGL2,* g*AXIN1,* g*DKK1,* g*COL2A1,* g*TWIST2,* g*RUNX2,* g*FAT2* showed a downward tread. The expression levels of g*WNT5A,* g*ROR2,* g*BMP4,* g*MSX1* were not affected (black text). Statistical analysis was done with one-way ANOVA followed by Dunnett's multiple comparison test. p value *<0.05, **= p< 0.01. Key: A-E: p<0.05 - red text, not significant - black text, genes show a downward trend – green text.

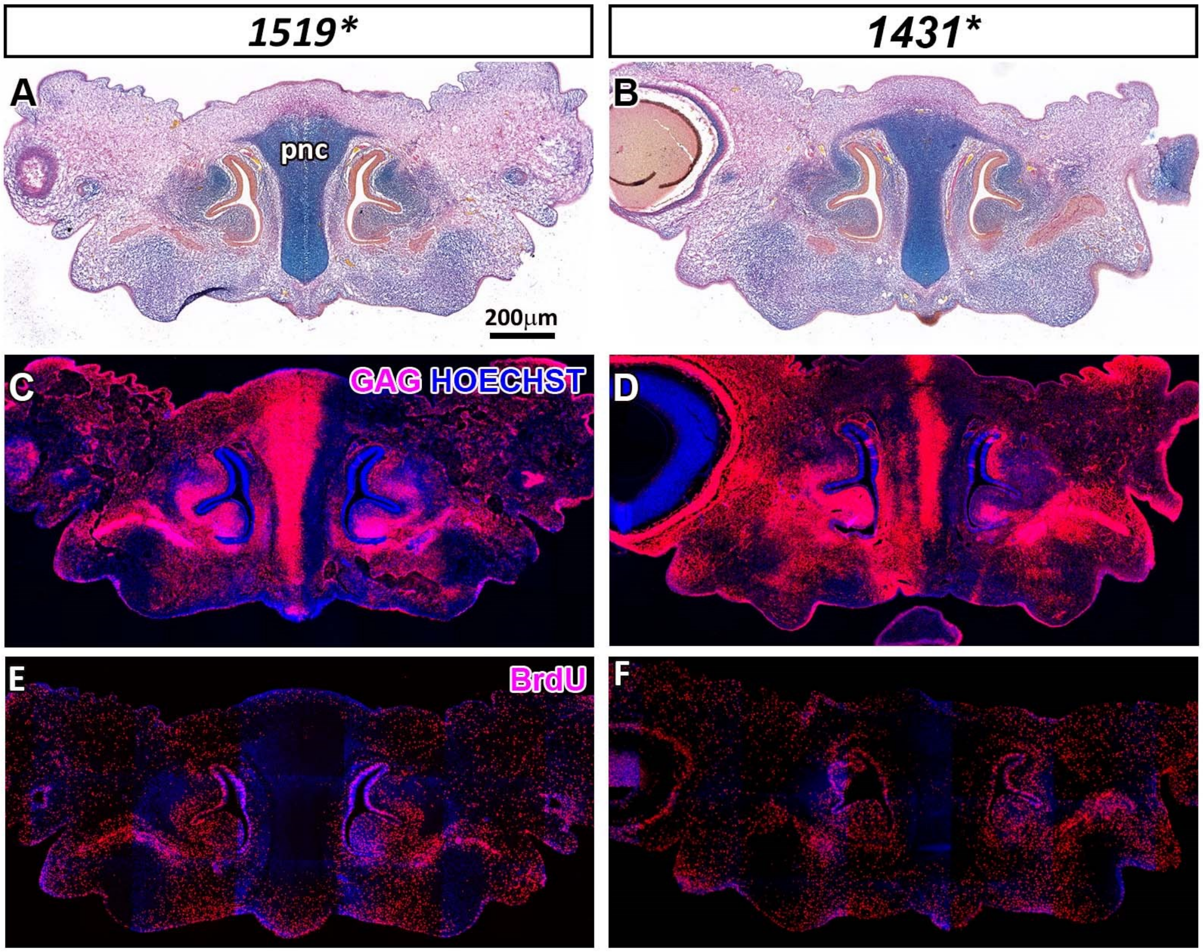


**Figure S4: hDVL1$^{1519*}$ and hDVL1$^{1431*}$ immunostaining at stage 29, 96h post injection**

Embryos injected into the right frontonasal mass with RCAS viruses containing h*DVL1*$^{1519*}$ and h*DVL1*$^{1431*}$ were fixed in 4% PFA 96h post-injection (HH29), embedded in paraffin and sectioned, 7μm sections. (A, B) Histological sections stained with alcian blue (stains cartilage) and picrosirius red (stains collagen fibers) show normal prenasal cartilage in the frontonasal mass. (C, D) Adjacent sections were stained with anti-GAG antibody (DSHB #GAG-pro 3C2-AMV) and counterstained with Hoechst (10μg/ml) to locate the virus in the tissues. Abundant viral spread was observed in all specimens injected with h*DVL1*$^{1519*}$ and h*DVL1*$^{1431*}$, n=3. (E, F) Embryos were treated with 10mM BrdU for 1 hour prior to fixation in 4% PFA. Anti-BrdU antibody (DSHB, #G3G4) was used to identify proliferating cells. Proportion of BrdU-positive cells did not show visible differences (data not quantified). Scale bar in A-F = 200μm Key: pnc – prenasal cartilage

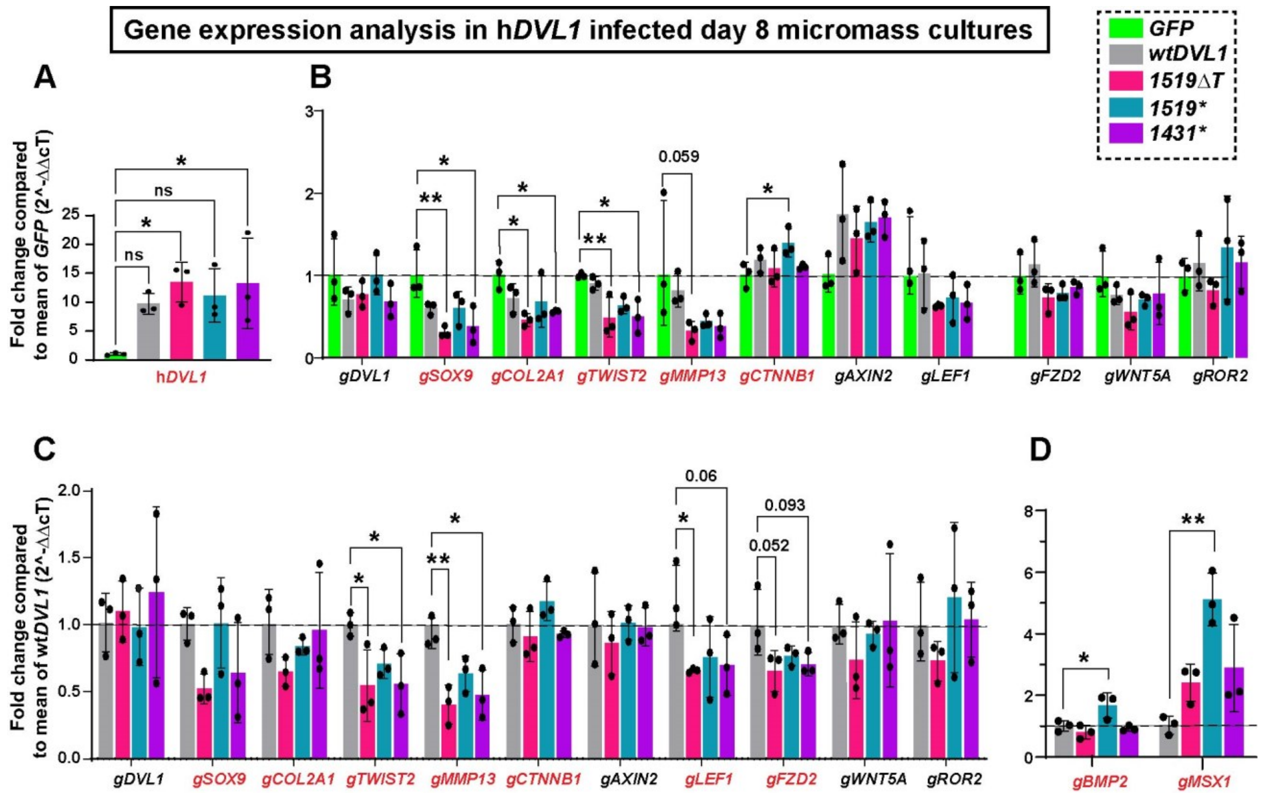


**Figure S5 Expression changes in WNT pathway genes in 8-day cultures infected with hDVL1 viruses qRT-PCR on frontonasal mass micromass cultures showed fold-change induction of hDVL1 (10-15-fold) in cultures treated with hDVL1 viruses compared to mean of GFP virus [2^(-ΔΔCt)] relative to the average levels of expression of the gene in GFP or wild-type hDVL1-infected cultures. Expression is normalized to 18s RNA. Three biological replicates with 8 cultures per biological replicate per virus type. The level of expression of human DVL1 was measured with human-specific primers. (A)There was a 10 to 15-fold higher level of hDVL1 all DVL1 virus infected culture conditions. (B) The RNA levels of gDVL1 remained unaffected after overexpression of wild-type or mutant DVL1 viruses. There is a significant decrease in levels of gCOL2A1 and gSOX9 in cultures infected with DVL1$^{1519\Delta T}$ and DVL1$^{1431*}$ viruses compared to GFP. Significant increase in expression of gCTNNB1 were caused by DVL1$^{1519*}$ compared to GFP. (C) The expression levels of gTWIST2, gMMP13, gLEF1, gFZD2 were lower in DVL1$^{1519\Delta T}$ and DVL1$^{1431*}$ virus infected cultures compared to wild-type DVL1. (D) BMP pathway genes gBMP2 and gMSX1 were upregulated by DVL1$^{1519*}$ compared to wild-type DVL1. Statistics done with one-way ANOVA followed by Dunnett's multiple comparison test using Prism 10.1.0. Key: *= $p<0.05$, **= $p< 0.01$, ns – not significant. Key: A-D: $p<0.05$ - red text, not significant -**

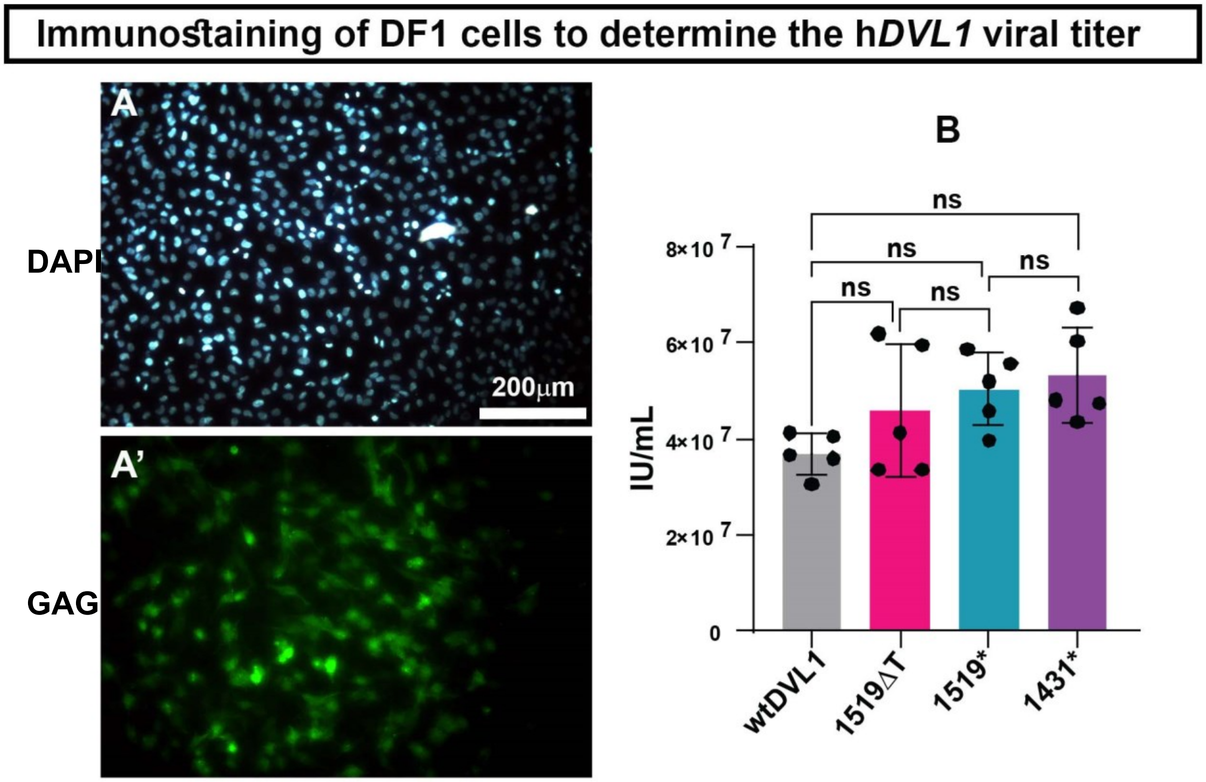


**Figure S6: Immunostaining on DF1 cells infected with high titer hDVL1 virus culture and quantification of viral titer**

(A, A’) Immunocytochemistry performed on DF1 chicken fibroblasts infected with 2µl of high titre viral stock were fixed at 36h post-infection. The cells were stained with anti-GAG antibody (virus) and counterstained with Hoechst. (B) Quantification of viral titer showed that all h*DVL1* viruses had higher than the recommended titer of 1 x $10^7$IU/mL for performing overexpression studies. Scale bar = 200µm. Key: IU – Infectious units, ns – not significant.

### Supplementary tables

#### Table S1: List of Robinow syndrome causing genes

| Gene/ OMIM ID | Function | Inheritance | Mutation type | OMIM entry |
|---|---|---|---|---|
| ***WNT5A*** **164975** | Ligand | Autosomal dominant | Missense | 180700 |
| ***DVL1*** **601365** | Cytoplasmic adaptor protein | Autosomal dominant | -1 frameshift | 616331 |
| ***DVL2*** **602151** | Cytoplasmic adaptor protein | Autosomal dominant | +1 frameshift | No entry |
| ***DVL3*** **601368** | Cytoplasmic adaptor protein | Autosomal dominant | -1 frameshift | 616894 |
| ***FZD2*** **600667** | Receptor- | Autosomal dominant | Missense, nonsense | No entry |
| ***ROR2*** **602337** | Receptor | Autosomal recessive | Missense, nonsense | 268310 |
| ***NXN*** **612895** | Stabilizer of *DVL* | Autosomal recessive | Deletion, missense | No entry |
| | | | | (Zhang et al., 2022) |

**Table S2: Knockout mouse models of Dvl isoforms**

| Genotype | Phenotype | Reference |
|---|---|---|
| ***Dvl1***$^{-/-}$ | Viable, fertile, no skeletal defects, social abnormalities, sensorimotor gating defects | (Lijam et al., 1997) |
| ***Dvl2***$^{-/-}$ | 50% die after birth, Cardiac defects, vertebral and rib malformations, spina bifida | (Hamblet et al., 2002) |
| ***Dvl3***$^{-/-}$ | Cardiac defects, abnormal orientation of hair cells, cochlear defects | (Etheridge et al., 2008) |
| ***Dvl1***$^{-/-}$ / ***Dvl2***$^{-/-}$ | Severe cardiac and skeletal defects, open neural tube, exencephaly, shortening of cochlear ducts, abnormal inner ear hair cell orientation | (Hamblet et al., 2002; Wang et al., 2006) |
| ***Dvl1***$^{-/-}$ / ***Dvl3***$^{-/-}$ | Mutants die between E13.5 - E15.5, cardiac defects | (Etheridge et al., 2008) |
| ***Dvl2***$^{+/-}$ / ***Dvl3***$^{-/-}$ | Abnormal head shape, forebrain defects, truncated snout, short and kinked trail, craniorachischisis, cardiac defects | (Etheridge et al., 2008) |
| ***Dvl2***$^{-/-}$ / ***Dvl3***$^{-/-}$ | Gastrulation defects, severe heart defects, posterior truncation lethality at E8.5 | (Etheridge et al., 2008) |

**Table S3: Patients carrying ADRS-DVL1 mutations reported in the literature**

| Total ADRS-*DVL1* patients | Male/Female | Clinical phenotype | Type of mutation (Frameshift) |
|---|---|---|---|
| **23** | 11/12 | Frontal bossing 17/19, Midface hypoplasia 22/23 Hypertelorism 21/23 | c.1496_1508del, c.1505_1517del, c.1508del, c.1518del, **c.1519del**, c.1529del, c.1556del, c.1570_1571del, c.1608_1623del, c.1612_1615del, c.1615del, c.1623del, c.1619_1631del |

(Zhang et al., 2022)(Hu et al., 2022)

**Table S4: Restriction free primer sequences used for cloning hDVL1 variants**

| ***DVL1* constructs** | **Primer sequence (5’→ 3’)** |
|---|---|
| ***DVL1***$^{1519\Delta T}$ | `Fw: GCTGCCCCCGGCCTCTGGGTCAG` |
| | `Rv: CTGACCCAGAGGCCGGGGGCAGC` |
| ***DVL1***$^{1519*}$ | `Fw: GCTGCCCCCTAACCTCTGGGTCAG` |
| | `Rv: CTGACCCAGAGGTTAGGGGGCAGC` |
| ***DVL1***$^{1431*}$ | `Fw: CTACGTCTTCGGGGATCTCTGCAGCTAAAATCTCG` |
| | `Rv: GTTGAGGTTCAGGGTGGCGAGATTTTAGCTGCAG` |

**Table S5: Analysis of wholemount embryos injected with DVL1 viruses**

| Virus type | Total embryos injected | % survival | Stage of collection | Total specimens (n) | Upper beak phenotype n, (%) | Premaxilla absent (n) |
|---|---|---|---|---|---|---|
| ***GFP*** | 35 | 69% | HH38 | 24 | 0 (0%) | 0 |
| **wt h*DVL1*** | 24 | 51% | HH38 | 14 | 8 (57%) | 8 |
| ***DVL1***$^{1519\Delta T}$ | 13 | 57% | HH38 | 13 | 12 (92%) | 12 |
| ***DVL1***$^{1519*}$ | 25 | 58% | HH38 | 13 | 0 (0%) | 0 |
| ***DVL1***$^{1431*}$ | 25 | 16% | HH38 | 4 | 1 (25%) | 1 |
| ***DVL1***$^{1431*}$ | 14 | 42% | HH33 | 6 | 4 (66%) | - |

### Table S6: Embryo stage and fixative post-injection

| Stage analyzed | Sample size (n) | Fixative | Experiment |
|---|---|---|---|
| **HH29** | 4-6 | 4% PFA | Histology, immunostaining |
| | 3 biologicals replicates | Flash frozen | qRT-PCR |
| **HH33** | 14 | 10% Formaldehyde | Phenotypic analysis |
| **HH38** | 13-35 | 100% ethanol | Skeletal staining |

### Table S7: Micromass cultures used for histology and immunostaining analysis

| Virus type | Total cultures collected for histology and immunostaining | |
|---|---|---|
| | **Day 6** | **Day 8** |
| ***GFP*** | 3 | 4 |
| **wt h*DVL1*** | 3 | 4 |
| ***DVL1***$^{1519\Delta T}$ | 3 | 6 |
| ***DVL1***$^{1519*}$ | - | 5 |
| ***DVL1***$^{1431*}$ | - | 5 |

### Table S8: Antibodies and immunofluorescence treatments

| Antigen Retrieval (steam 15 minutes, 95°C) | Permeabilization/ Pre-treatment | Block Goat serum | Primary antibody, source, dilution (Overnight, 4°C) | Secondary Antibody and counterstain |
|---|---|---|---|---|
| **10mM Sodium Citrate** | 0.1% Triton X-100 in 1XPBS for 15 min | 10% Goat serum (Sigma G9023), 0.1% tween-20 in 1X PBS, (90 minutes, RT) | **Ctnnb1/PY489-ß-catenin** Developmental Studies Hybridoma bank (DSHB), 5µg/ml, Mouse monoclonal | ThermoFisher, Cy5 goat anti-mouse, #A10524,<br><br>ThermoFisher, Cy5 goat anti-rabbit, #A10523,<br><br>ThermoFisher, Alexa Fluor™ 488 goat anti-mouse, A11029,<br><br>ThermoFisher, Alexa Fluor™ 488 goat anti-rabbit A11034, 1:200 at RT in dark, 90mins<br><br>**Hoechst** 10µg/ml (Sigma 33568) in 1 X PBS (RT in dark for 30 min) |
| **10mM sodium citrate** | 0.5% hyaluronidase in Hank's Balanced Salt Solution, 45min (following antigen retrieval) | | **Collagen type II** DSHB, #II-II6B3, 5µg/ml, Mouse monoclonal | |
| **10mM Sodium Citrate** | 0.5% hyaluronidase in Hank's Balanced Salt Solution, 45min (following antigen retrieval) | | **Collagen type II** Proteintech®, 28459-1-AP, 1:50, Rabbit polyclonal | |
| **1X Diva Decloaker (BioCare Medical)** | 0.1% Triton X-100 in 1XPBS for 15 min | | **Gag-pro AMV-3C2** DSHB, 1:4, Mouse monoclonal | |
| **10mM Sodium Citrate/ 1X Diva Decloaker (BioCare Medical)** | 0.1% Triton X-100 in 1XPBS for 15 min | | **SOX9** Sigma-Aldrich, #HPA001758 1:200, Rabbit Polyclonal | |
| **10mM Sodium Citrate/ 1X Diva Decloaker (BioCare Medical)** | 0.1% Triton X-100 in 1XPBS for 15 min | | **TWIST1** Twist2C1a Abcam #ab50887 1:50, Mouse monoclonal | |
| **0.3% Triton X-100 in 1XPBS for 15 min** | 0.1% Triton X-100 in 1XPBS for 15 min | | **Flag (DYKDDDDK)** Thermo Fisher PA1-984B, 1µg/ml, Rabbit Polyclonal | |
| **1X Diva Decloaker (BioCare Medical)** | 0.1% Triton X-100 in 1XPBS for 15 min | | **BrdU** DSHB, G3G4, 5µg/ml, Mouse monoclonal | |

**Table S9: Quantitative-PCR primers**

| Gene | Accession # | Forward Primer | Reverse Primer |
|---|---|---|---|
| *hDVL1* | NM004421.3 | CAGCATAACCGACTCCACC | TGATGCCCAGAAAGTGATGTC |
| *gDVL1* | XM015297120.1 | CTCCCATTGAGAGGACAGGT | TGTTTCGTTGTCCAGTCCAT |
| *gWNT5A* | NM003392 | CAATGGCTTCTCAGTACCTCG | ACATCTGCACAGGGTTCATG |
| *gCTNNB1* | NM 205081 | CTTGGACTTGACATTGGTGC | CAGAGTGGAAAGAACGGTAGC |
| *gROR2* | NM 001080716.1 | AGTGCTGGAATGAATTTCCC | GGGAACTTGTTTGTGTGGTG |
| *gFZD2* | NM204222.2 | CTCCCATTGAGAGGACAGGT | TGTTTCGTTGTCCAGTCCAT |
| *gLRP5* | NM001012897 | ACCAAAGCCAGAACCCAG | CAGCACCATCCCTATTGACTC |
| *gATF2* | NM 204904 | GCCAGCGTTTTACCAATGAG | AGTTGGTGTTGGTGTCTGATC |
| *gLEF1* | NM_001398055.1 | CATCAAGTCCTCGCTGGTC | GCCCTTGTCATGGTAGGAATC |
| *gCOL2A1* | NM 204426.1 | GGACCAGCAAGACGAAAGAC | CGTAGCTGAAGTGGAAACCG |
| *gSOX9* | NM_204281.2 | CTGGGCAAGCTGTGGAG | GGTTGGTACTTGTAGTCGGG |
| *gTWIST1* | NM_204739 | GACTCCAAGATGGCAAGCTG | CTCCATTCTCCACACCGAGA |
| *gTWIST2* | NM_204679 | GAGTTATGCCTTCTCAGTCTGG | ACGTCCCAATTCCACTTCAG |
| *gBMP2* | NM204358 | GCTGTTTTGAGGTGGATTGC | AGGCACTGTTCTCTTTGTCC |
| *gBMP7* | XM 417496 | GTCAAACATCGCAGAGAACAG | TCCTTCACAGTAATACGCAGC |
| *gNOGGIN* | NM204123 | ACTTTATGGCTATGTCCCTGC | AACTCCAGCCCCTTGATTTC |
| *gBMPER* | NM 001007080 | AGCTGTCCTCATGGTAAAATCC | AACGTACTGACATGTTCCCTG |
| *gSP7* | XM015300329.3 | GTTCGTCTGCAATTGGCTCT | AATTTCTTCTCGCGGGTGTG |
| *gRUNX2* | NM204128 | ACCTAGTTTGTTCCCTGAACG | GTAATCTGACTCTGTCCTTGTGG |
| *gMSX1* | NM_204559 | TTCGGTCAAATCGGAGAACTC | TTCGTCTTGTGCTTCCTCAG |
